# New Insights into Microbial Nitrogen Utilization in the Rumen Enabled by Genome-Resolved Multi-Omics

**DOI:** 10.1101/2025.03.23.644819

**Authors:** Ming Yan, Jeffrey Firkins, Jiarong Guo, Alejandro Relling, Zhongtang Yu

## Abstract

Optimizing nitrogen (N) utilization in ruminant production systems holds both economic and environmental significance. However, traditional paradigms of N metabolism, derived primarily from well-studied model rumen bacteria, cannot fully capture the diverse and complex N metabolic dynamics within the rumen ecosystem. To address this gap, we utilized comparative genomics and genome-resolved multi-omics analyses using a curated set of microbial genomes to investigate N assimilation and regulation in rumen microbes. We discovered that canonical mechanism of ammonia assimilation and regulation, such as the glutamine synthetase (GS)/glutamate synthase (GOGAT) pathways and its regulatory proteins, are absent in the genomes of many key and predominant rumen microbes, which likely utilize alternative pathways for ammonia assimilation. These findings challenge the applicability of *E. coli*-based N regulation models to rumen bacteria. We further linked polysaccharide utilization and ammonia assimilation across hundreds of rumen microbial species. Furthermore, we identified specific microbial species involved in ureolysis and denitrification, as well as phages carrying auxiliary metabolic genes that perform N assimilation. Using an animal trial involving 11 pairs of lamb twins in a crossover design, we demonstrated that dietary crude protein (CP) concentrations had minimal impact on rumen microbiome composition and expression of N assimilation genes. Instead, shifts in concentrate levels triggered alterations in N assimilation, including increased expression of amino acid biosynthesis pathways. These findings indicate a nuanced, species-specific microbial response to dietary interventions, highlighting the limitations of traditional N metabolism models applied to rumen microbes and the need for more granular studies of rumen microbial ecosystems.

## INTRODUCTION

Efficient utilization of dietary nitrogen (N) is critical for sustainable ruminant production. Microbial N contributes up to 80% of the total absorbable protein in the intestine of ruminants [1, 2], yet N utilization efficiency (NUE) in ruminants remains low, with over 70% of ingested N excreted in manure [3]. This inefficiency not only reduces profitability due to the high cost of dietary N [4] but also raises multiple environmental concerns, such as greenhouse gas (GHG) emissions (e.g., N_2_O, which is 298 times more potent than CO_2_ [5] and contributing to PM2.5), soil acidification, and water pollution [6].

Decades of research to improve NUE in ruminants, including strategies such as N and energy synchronization via dietary formulation, have yielded inconsistent and often marginal outcomes [7, 8]. These limited gains underscore a critical gap in our understanding of the rumen microbiome as a dynamic, interacting community that encompasses a diverse array of bacteria with distinct metabolic strategies and ecological niches. While most rumen bacteria participate in dietary protein degradation and ammonia assimilation, they differ in substrate preferences, metabolic capacities, and ecological niche partitioning, each with distinct implications for N utilization. For example, ureolytic bacteria hydrolyze urea to ammonia, playing a direct role in urea recycling and urine excretion, while hyper-ammonia-producing bacteria (HAB) rapidly ferment amino acids and peptides, leading to excessive deamination in the rumen [9]. Rumen phages further influence N metabolism in this ecosystem by modulating microbial N processing and utilization, either by augmenting microbial metabolism capacities with phage-encoded auxiliary metabolism genes (AMGs) or by host lysis, thereby contributing to intra-ruminal N recycling [10].

Traditional culture-dependent methods have facilitated the identification of certain bacterial species involved in protein degradation and deamination within the rumen [11]. However, the inherent phenotypic bias associated with culture-dependent methods and the limited availability of cultured rumen bacteria limit the extrapolation *in vitro* metabolic observation of cultured rumen bacteria to the *in vivo* rumen ecosystem. Recent advances in genome-resolved metagenomics have greatly expanded our understanding of the phylogenetic and functional diversity of complex microbial ecosystems, including the rumen microbiome, by uncovering the metabolic pathways of previously uncultivated bacteria and by enabling comparative genomic analyses, along with other omics technologies like metatranscriptomics [12–14]. These tools offer unprecedented opportunities to systematically investigate the N utilization strategies of rumen bacteria across different niches. To address this, we employed a multi-omics approach to investigate microbial N utilization in the rumen. We first performed a comparative genomic analysis of thousands of rumen microbial genomes to systematically characterize their N metabolism strategies. Subsequently, we performed an animal trial with a split-plot crossover design involving 11 pairs of lamb twins, fed diets with varying concentrate-to-forage ratios and crude protein (CP) concentrations. By integrating metagenomic and metatranscriptomic data, we aimed to provide a genome-resolved understanding of how the rumen microbiomes regulate N utilization under different dietary regimes. This integrative approach enhances our understanding of the N metabolic capacities and interactions of rumen bacteria, with implications for improving NUE in ruminants.

## MATERIALS AND METHODS

### Animal trial and sampling

All procedures involving animals were approved by The Ohio State University Institutional Animal Care and Use Committee (protocol number: 2020A00000080-R1). Eleven pairs (12 pairs initially, one pair removed due to a health issue) of lamb twins were allocated to four dietary treatments in a split-plot crossover design, fed diets with varying concentrate-to-forage ratios (as the main-plot treatment, assigned to each pair of twins) and CP concentrations (as split-plot treatment, within each pair of twins). Specifically, one twin of each twin pair was either assigned to a high-forage diet (70% forage) or a low-forage diet (30% forage). Within each twin pair, one lamb was assigned to a low CP diet (10% of dry matter) in period one and then switched to a high CP diet (13% of dry matter) in period two, while the other lamb followed the reverse dietary sequence (Fig. S1; also see Table S1 for the detailed nutrient composition of the diets). The low- and high-CP diets were formulated to be isoenergetic at the net energy for maintenance level for each forage group, following the model outlined in the National Research Council (2007) [15]. The washout period between periods 1 and 2 lasted 14 days, during which a blended diet comprising an equal amount from periods 1 and 2 was provided. Animals had *ad libitum* access to fresh water throughout the study. On the last day of each feeding period, rumen fluid samples were collected from each lamb via oral tubing 2 h after the morning feeding following the procedure described previously [16]. Specifically, about 10 mL of each rumen fluid was discarded to avoid saliva contamination before collecting a rumen fluid sample from each lamb. Four mL of each rumen fluid sample was mixed with 0.8 mL of meta-phosphoric acid (25% (w/v), made fresh) for volatile fatty acids (VFA) analysis; 2.5 mL was mixed with 150 µL of HCl (5N, freshly made) for ammonia analysis; 3 mL was mixed with 10 mL of RNALater® for RNA isolation; and 3 mL was collected for DNA extraction. During collection, all rumen fluid samples were snap-frozen in dry ice on-site and then stored at −80 °C in the lab until analyses. Ammonia concentrations were measured using the colorimetric method described by Chaney and Marbach [17], while VFA concentrations were determined via gas chromatography [18].

### DNA sequencing and metagenomic analyses

DNA was extracted using the repeated bead-beating plus column purification method described by Yu and Morrison [19]. Metagenomic sequencing was performed on the DNBSEQ platform (BGI, Beijing, China) with paired-end reads of 150 bp (PE150) at Innomics Inc. (Sunnyvale, CA). Default parameters were used for each bioinformatic tool unless otherwise specified. The raw reads were quality-controlled with fastp [20] to remove low-quality reads. Potential host-derived reads were filtered out using Bowtie2 [21]. The resulting clean reads were assembled both individually and through co-assembly for each animal or treatment with MEGAHIT [22] with default parameters. The contigs from the three approaches were binned using MaxBin2 [23], MetaBAT2 [24], and CONCOCT [25]. The final refined bin sets were subsequently generated using the DAS Tool [26]. The completeness and contamination of the three sets of refined bins were assessed using CheckM2 [27]. Among the three assembly approaches, co-assembly per animal yielded the highest number of high-quality metagenome-assembled genomes (MAGs). These MAGs were dereplicated using dRep [28] at a 95% ANI threshold. The dereplicated MAGs with >50% completeness and <10% contamination were retained for downstream analyses.

We also collected additional MAGs (7,176 MAGs described by Yan and Yu [29] and 4,941 MAGs described by Stewart, et al. [30]) and genomes from the Hungate1000 rumen microbial genome collection [31] to expand the collection of rumen MAGs and genomes. These MAGs and genomes were checked for quality using CheckM2. The high-quality MAGs (> 90% completeness, < 5% contamination) were combined with those derived from the lamb rumen samples and dereplicated at 98% ANI to preserve multiple MAGs per species, resulting in a non-redundant collection of 5,479 MAGs/genomes. The taxonomy of these MAGs was assigned using GTDB-Tk2 [32]. To ensure robust representation for downstream analyses, species represented by fewer than 2 MAGs were excluded, yielding 2,384 bacterial MAGs/genomes representing 741 species across 263 genera. This curated dataset of 2,384 MAGs, referred to as Rumen Microbial MAGs hereafter, was used for comparative genomic analyses.

### Comparative genomics analysis

To comprehensively understand the N utilization pathways within the rumen, we annotated all the rumen microbial MAGs for genes involved in N utilization (detailed in Table S2). For a function to be considered encoded within a genome, all constituent subunits of the associated gene cluster must be identified. For instance, *ure*C, *ure*B, *ure*A, *ure*E, *ure*F, *ure*G, *ure*D, and *ure*H must be present to confirm that the genome encodes urease. Because we focused on species represented by multiple MAGs to ensure robust analyses, we considered a gene as present in a species only when at least 80% of the MAGs encoded the same genes (to avoid misidentification of a gene in a species due to assembly or binning errors). A gene was considered absent in a species when it was absent in all of the MAGs of the same species. We retained species only if all N-related genes had consistent annotation (present or absent) across the MAGs within the species to build the phylogenetic trees. Specifically, the sequences of the concatenated marker genes specified in GTDB for bacterial genomes encoding various pathways for ammonia assimilation (472 species) or assimilation regulation (192 species) were aligned using GTDB-Tk2 (gtdbtk align). Phylogenetic trees were constructed from these alignments using IQ-TREE (option: -redo -bb 1000 -m MFP -mset WAG,LG,JTT,Dayhoff -mrate E,I,G,I + G -mfreq FU -wbtl). The trees were visualized using iTOL [33].

### Taxonomic profiling and functional annotations of the MAGs reconstructed from the rumen microbiome of lambs

Taxonomic profiling of the rumen microbiome was performed using the Kraken2 [34] classifier and the GTDB taxonomy [35]. The Kraken2 output was then used to generate abundance profiles at the genus and species levels using Bracken [36]. Only taxa with a relative abundance ≥0.001% in more than 50% of the samples were retained for downstream statistical analysis. The distribution of the MAGs across the samples was analyzed using CoverM (https://github.com/wwood/CoverM) with the default setting. Functional annotation of the reconstructed MAGs was conducted using DRAM [37]. Open reading frames (ORFs) of these annotated MAGs were concatenated into a catalog, which served as a reference for subsequent mapping metatranscriptomic reads in genome-resolved analyses.

### Identification of phages and AMG predictions

Phage sequences were identified from the assembled contigs longer than 5 kb using both VirSorter2 [38] and VIBRANT [39], as described in our previous studies [29, 40]. The quality of the identified phage sequences was assessed using CheckV [41]. Only the identified phage genomes with medium or higher quality (50-90% completeness, based on CheckV results) were subjected to AMG identification using DRAM-v [37], with a focus on a set of curated genes (Table S2) using the criteria established by Yan, et al. [40]. All phages harboring putative AMGs were manually curated to confirm accurate demarcation of host-phage genome boundaries and phage sequence integrity.

### RNA extraction, sequencing, and metatranscriptomics analyses

Following the manufacturer’s protocol, total RNA was isolated from the rumen fluid samples using the Quick-RNA Miniprep Plus Kit (Zymo Research). To enrich for microbial mRNA, ribosomal RNA was depleted with the NEBNext rRNA Depletion Kits (Bacterial and archaeal rRNA depletion; New England Labs, Ipswich, MA). Sequencing libraries were then prepared using the Rapid Directional RNA-Seq Kit 2.0 (PerkinElmer, Waltham, MA) and sequenced on the NovaSeq X Plus Sequencing System (Illumina, San Diego, CA, USA) with 2 x 150 bp paired-end reads using the NovaSeq X Series 1.5B Reagent Kit (300 cycles). Quality control of raw reads was performed with fastp, and residual rRNA sequences were removed using SortMeRNA [42]. The resultant mRNA reads were mapped to the catalog of ORFs predicted from the Rumen Microbial MAGs using RSEM [43] to quantify gene expression profiles for each MAG. The read-based transcriptional profiles were further analyzed at both gene and pathway levels using HUMAnN3 [44], enabling the identification of key functional pathways associated with N metabolism.

### Statistical analysis

All statistical analyses were performed in R. Specifically, rumen fermentation profile data were analyzed with a linear mixed-effect model implemented in the lme4 package. In this model, forage and CP concentrations and their interaction were treated as fixed effects, while twin pairs were included as random effects, with individual lambs nested within each twin pair. Microbial abundance and transcriptional profiles were analyzed using LinDA [45]. For differential abundance analyses, *p*-values were adjusted for multiple comparisons using the Benjamini-Hochberg (BH) procedure to control false discovery rate (FDR). MAG-based analyses were restricted to prevalent MAGs detected in >80% of the samples within both forage levels. For MAGs present in >80% of the samples from only one forage level, a paired Wilcoxon test (paired by CP concentration) was conducted, since the effects of forage level, its interaction with CP concentration, and the random effect of twin pairs and individual lambs were not considered relevant. Statistical significance was declared at *p* < 0.05 for fermentation profiles and an FDR-adjusted *q* < 0.05 for microbial abundance and transcriptional profile data.

## RESULTS AND DISCUSSION

### Rumen bacteria employ species-specific strategies for polysaccharide degradation and N assimilation

Using thousands of recently sequenced rumen microbial genomes, we performed systematic annotation to investigate the pathways involved in N and polysaccharide utilization (Fig. 1). Among the 472 bacterial species examined, the majority lacked a complete glutamine synthetase-glutamate synthase (GS-GOGAT) pathway, as evidenced by the absence of GOGAT in their genomes. This finding suggests that most rumen bacteria likely use glutamate dehydrogenase (GDH) for ammonia assimilation. Additionally, alanine dehydrogenase likely plays a significant role in ammonia assimilation, particularly at high ammonia concentrations. This is consistent with reports identifying alanine as the predominant amino acid at high ammonia conditions within the rumen [46] and increased alanine dehydrogenase activity in response to elevated ammonia levels [47]. While alternative pathways, such as aspartase, may also participate in ammonia assimilation, further research is required to determine whether their roles are assimilatory or dissimilatory. The identification of symbiotic and metabolically specialized bacteria underscores the complexity of N metabolism within the rumen ecosystem.

**Fig. 1:**
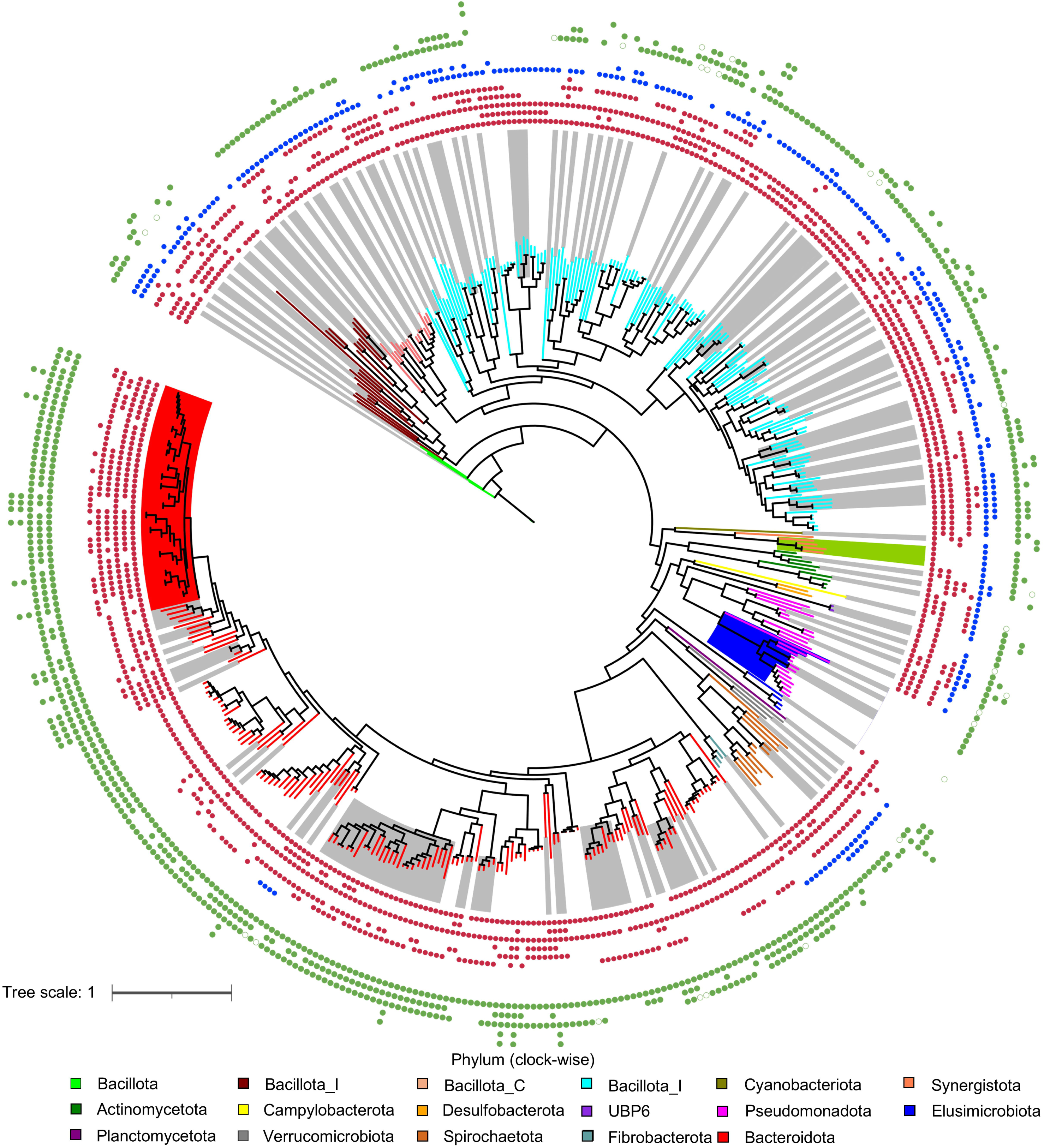
Overview of N utilization-related genes encoded by rumen bacteria and their glycoside hydrolases. The data represent the N assimilation genes encoded by 472 genomes of individual rumen bacterial species. Outer rings (in green, from innermost to outermost) represent the detected glycoside hydrolases capable of hydrolyzing chitin, pectin, starch, cellulose, and hemicellulose (green hollow circles indicate annotation inconsistent within the species). Middle rings (from innermost to outermost) represent the detected glutamine synthetase, glutamate synthase, glutamate dehydrogenase, alanine dehydrogenase, asparagine synthetase, aspartase, amino acid transporter, and peptide transporter. The genera of *CAG-267*, *CAZU01*, *JAFUYI01*, *MGBC133411*, and *RUG410* within the phylum *Pseudomonadota* that do not encode any of the ammonia assimilation genes are highlighted in blue. The genus *Prevotella* is highlighted in red, and the genus *JAFUXM01* is highlighted in green. Each leaf represents an individual species, with its color indicating the corresponding phylum to which the species belongs. The branch within each color belongs to the same genus (white or grey). A gene (or gene cluster) is considered present in a species only when at least 80% of the genomes within that species carry the gene (or gene cluster). A gene (or gene cluster) is considered absent in a species when the gene is absent in all of the genomes within that species (or gene cluster).

Notably, some rumen bacteria rely exclusively on amino acids to meet their N requirement. For example, the genus *JAFUXM01* (highlighted in green in Fig. 1) encodes amino acid transporters but exhibits an incomplete GS-GOGAT pathway and lacks GDH. In contrast, the genomes of several bacterial orders of the class Alphaproteobacteria, including RF32 and Rs-D84, lack genes associated with ammonia assimilation (highlighted in blue in Fig. 1) or amino acid or peptide transports. These bacteria likely have reduced metabolic capacity with a potential symbiotic lifestyle, as recently observed in the MAGs from fecal microbiomes of farm animals (camels, yaks, and cows) [48]. Further investigation is warranted to elucidate these bacterial symbiotic relationships and ecological roles in the rumen ecosystem. Moreover, N assimilation strategies are not conserved across species within genera. For instance, within the genus *Prevotella* (highlighted in red in Fig. 1), certain species employ both GS-GOGAT and GDH for N assimilation, whereas others utilize GDH alongside other enzymes like aspartase. These findings suggest that N assimilation strategies within the rumen ecosystem are highly species-specific, reflecting their ecological roles and evolutionary adaptations. These findings also align with observed variations in rumen microbiome composition and NUE in ruminants [49].

The regulatory mechanisms of N assimilation in rumen bacteria also differ from the well-established paradigm observed in model organisms like *E. coli*, which involves the PII protein, glutamine synthetase, adenylyltransferase, PII uridylyl-transferase, and a two-component system NtrB/NtrC (also NRII/NRI). Genomic analysis of 192 rumen bacterial species across nine phyla revealed the absence of the complete N assimilation regulatory system of *E. coli* (Fig. S2). Similar observations have been reported in Bacteroidetes by Firkins and Mackie [50]. The absence of these systems suggests that ammonia availability is not a limiting factor in the rumen, possibly rendering these regulatory mechanisms unnecessary. Indeed, transcriptomic comparisons of wild-type and *gln*G mutant *E. coli* monocolonizing the mouse gut revealed minimal differences, with most *ntr*C-regulated genes being downregulated when compared to the expression profiles of wild-type *E. coli* in laboratory cultures under N-limiting conditions [51]. This again suggests that N is not limiting for *E. coli* in the mouse gut, making the induction of NtrC-dependent genes unnecessary. Further supporting this notion, Kim, et al. [52] demonstrated a similar level of expression of the GS-GOGAT system in *Prevotella ruminicola* 23 between ammonia-limiting and non-limiting conditions. Collectively, these findings imply that the high ammonia availability resulting from rapid dietary protein degradation in the rumen might have led to the evolutionary loss of the ATP-dependent GS-GOGAT system and related regulatory genes in certain rumen bacteria.

Microbial protein synthesis requires both N sources and energy input. Previous studies have established a simplified model in which cellulolytic bacteria predominantly use ammonia as their primary N source, whereas amylolytic bacteria utilize a broader range of N sources, including ammonia, amino acids, and peptides [8]. However, studies have demonstrated that cellulolytic bacteria also utilize N sources other than ammonia. For example, *Fibrobacter succinogenes* S85, a specialized cellulolytic bacterium within the rumen [53], was shown to accumulate ^14^C-labelled peptides and amino acids [54]. Moreover, various cellulolytic bacteria utilize different amino acids to varying degrees [55, 56]. The influence of carbon sources adds complexity to the relationship, as cellulolytic bacteria exhibit increased growth on cellobiose but not cellulose in the presence of peptides [57]. Our comparative genomic analyses revealed that the genomes of most cellulolytic/xylanolytic bacteria, particularly those within the phylum Bacteroidota, encode cellulase and/or hemicellulase but lack amino acid or peptide transporter genes. Additionally, these bacteria encode amylases. Conversely, most species in the phyla Bacillota, Bacillota_I, Bacillota_A, and Bacillota_C encode amino acid and/or peptide transporters in their genomes, with some also carrying cellulase- and hemicellulase-coding genes. These findings highlight the diverse strategies employed by rumen bacteria for N and carbohydrate utilization (Table S3 and Table S4). Furthermore, they corroborate earlier culture-dependent studies demonstrating such metabolic differentiation and emphasize the need for species-resolved genomic analyses to accurately characterize the functional diversity of rumen bacteria and avoid mischaracterization inherent in genus-level metataxonomic analyses.

### Rumen bacteria and phages carry genes encoding specialized N utilization capacities

In addition to ammonia assimilation, ureolytic activity is important as it directly relates to urea recycling and N excretion. In this study, we identified nine rumen bacterial species whose genomes encode the complete urease gene clusters, which were grouped into four distinct types based on their architecture (Fig. 2a). Type III comprises those of *Succinivibrio sp000431835* and *Succinivibrio sp900315395*, while type IV was found in *Treponema_D sp002373205* and has an adjacent urea transporter gene cluster (*urt*A, *urt*B, *urt*C, *urt*D, and *urt*E). Types I and II lack urea transporter genes. A recent functional gene-guided enrichment study successfully isolated 12 ureolytic bacteria and found five distinct types of urease gene clusters [58]. However, neither the ureolytic bacteria nor types II, III, and IV of urease clusters identified in this study were represented among these isolates. These findings expand the taxonomic and genetic diversity of ureolytic bacteria in the rumen microbiome and underscore its ecological versatility, emphasizing its potential role in influencing N utilization.

**Fig. 2:**
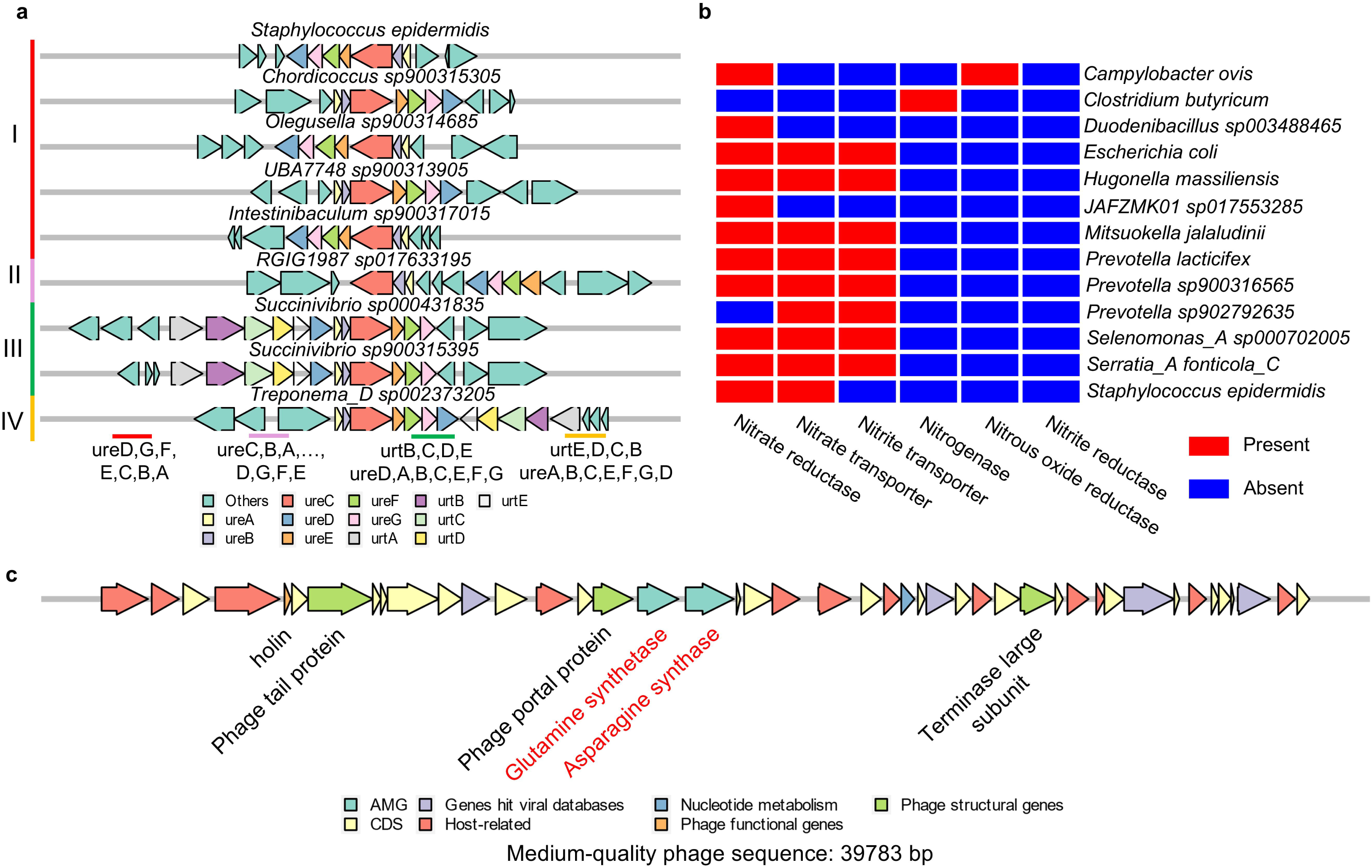
Identified rumen bacterial urease genes, denitrification genes, and species carrying these genes, along with one viral MAG carrying ammonia assimilation genes. **a**, Schematics of urease gene clusters identified among rumen ureolytic bacteria; **b**, Rumen bacteria identified to carry denitrification genes. Each species is represented by at least 2 genomes. A gene (or gene cluster) is considered present in a species when at least 80% of the genomes within that species carry this gene (or gene cluster). A gene (or gene cluster) is considered absent in a species when the gene is absent in all the genomes within that species (or gene cluster). **c**, A medium-quality (as determined by CheckV) viral MAG encoding glutamine synthetase and asparagine synthase as auxiliary metabolism genes.

Among the newly identified ureolytic species, all except *Olegusella sp900314685* encoded a high-affinity ammonium transporter (*amt*B). In contrast, only approximately half (250 out of 472) of the previously described species possess *amt*B (Table S4). Of these 250 species, only 80 lack genes encoding amino acid/peptide transporters, suggesting exclusive reliance on ammonia as their nitrogen source. A previous study has also observed the physical propinquity of *amt*B and *gln*K (which encodes an N regulatory protein) in the rumen microbial genomes [50]. Therefore, *amt*B and its regulatory mechanisms may function as an adaptive strategy for rumen microbes, enabling them to respond to fluctuating ammonia concentrations and compete for N under variable ammonia conditions.

We identified rumen bacteria whose genomes encode nitrate reductase and nitrate transporters. These include species of *Prevotella* and nine other rumen bacteria across different genera (Fig. 2b). Although both assimilatory (to ammonium) and dissimilatory (to N_2_) routes of nitrate reduction are observed in the rumen [59], the specific pathways used by these bacteria remain to be determined. In addition, we identified rumen bacterial species that carry genes encoding nitrogenous oxide reductase (e.g., *Campylobacter ovis*) and nitrogenase (i.e., *Clostridium butyricum*). While nitrate has been used to reduce methane production in the rumen, the supplementary nitrate is predominantly reduced to ammonia rather than to N_2_ through denitrification [60]. Similarly, N fixation in the rumen is minimal [61] and the nitrogenase activity of rumen bacteria was negligible [62, 63]. This aligns with the rarity of rumen bacteria that encode nitrogenase, only *Clostridium butyricum*, identified in the present study.

We identified a medium-quality (determined by CheckV) viral MAG (vMAG) encoding two ammonia assimilation-related AMGs: asparagine synthase and glutamine synthetase. While previous studies identified AMGs involved in N metabolism (e.g., denitrification and a PII protein that regulates ammonia assimilation) [40, 64], this is the first report of AMGs directly involved in ammonia assimilation. Although this vMAG is incomplete, the identified AMGs were flanked by multiple viral hallmark genes, indicating that the AMGs are not part of the host genome. Additionally, the identification of the holin gene and multiple other genes matching viral databases (VOGDB or RefSeq Viral) suggests that these AMGs are not part of a genomics island. These AMGs may enhance N assimilation in infected rumen bacteria, potentially supporting viral replication through increased protein and nucleotide synthesis. This aligns with previous observations of increased N assimilation in phage-infected bacteria [65]. The identification of N-related AMGs in phages suggests another mechanism by which viruses mediate N cycling within the rumen ecosystem.

### Rumen archaea likely utilize amino acids in addition to ammonia

We examined the N assimilation capacities of nine archaeal species across three archaeal phyla (Fig. S3), including both the well-studied phylum Methanobacteriota and less-explored phyla Thermoplasmatota (e.g., *Methanomethylophilus sp001481295* and *UBA71 sp015063165*) and Halobacteriota (e.g., *Methanomicrobium mobile*). Similar to the rumen bacteria, none of the examined archaeal species possesses GOGAT. Some archaea, such as *Methanomicrobium mobile*, likely use preformed amino acids to meet their N requirements, as evidenced by the lack of GDH but the presence of amino acid transporters. Additionally, none of the examined archaeal species contained proteins involved in the classic ammonia assimilation paradigm established in *E. coli*. These findings highlight a distinct N assimilation strategy in rumen archaea, with potential reliance on external amino acids and other N sources, underscoring their ecological specialization within the rumen ecosystem. Due to their low abundance, rumen archaea do not likely have a significant direct effect on NUE, but their N utilization strategies may inform the development of effective methane mitigations.

### Dietary forage and CP concentrations influence rumen fermentation profiles in lambs

Optimizing rumen fermentation efficiency is critical for improving nutrient utilization in ruminants, particularly when diets are formulated for varying forage and CP concentrations. Although previous studies have identified dominant microbial taxa under different dietary conditions, taxonomic resolution has largely been restricted to the genus level due to the limitations of metataxonomic or gene-centric metagenomic approaches. However, as discussed in previous sections, many rumen bacteria employ species-specific polysaccharide and N utilization strategies, as exemplified by species of the genus *Prevotella*. To better understand the nutritional roles of rumen bacteria, we investigated their responses to varying N and forage levels at a higher resolution by conducting a feeding trial involving 11 pairs of lamb twins fed diets with different forage and CP concentrations (Fig. S1), combined with genome-resolved metagenomics. We first measured the rumen fermentation profile (Table 1) and verified that neither the feeding period nor the crossover effect was significant. We further compared the treatment effects. As expected, the ammonia concentration was significantly greater in the high CP group (*p=*0.02), reflecting the increased deamination by rumen microbes. Additionally, propionate concentration was significantly higher in the low-forage group (*p<*0.01), aligning with the greater fermentation of non-fiber carbohydrates typical of such diets. A significant interaction was observed between CP and forage concentration for butyrate concentration (*p=*0.04), with the low forage and low CP group exhibiting significantly lower butyrate concentrations than the other dietary treatments. Dietary adjustments in forage and CP concentrations likely exert interactive effects on key rumen fermentation parameters, which may have downstream nutritional implications in lambs.

**Table 1:** Rumen fermentation profiles of lambs fed diets varying levels of forage and crude protein (CP).

| Rumen fermentation | Low Forage |  | High Forage |  | SEM | Effects (*P*-values) |  |  |
| --- | --- | --- | --- | --- | --- | --- | --- | --- |
|  | Low CP | High CP | Low CP | High CP |  | Forage-level | CP-level | Forage x CP interaction |
| Acetate (mmol/L) | 51.90 | 41.56 | 51.92 | 52.84 | 2.184 | 0.06 | 0.86 | 0.12 |
| Propionate (mmol/L) | 28.66<sup>A</sup> | 27.61<sup>A</sup> | 14.81<sup>B</sup> | 14.70<sup>B</sup> | 1.378 | <0.01 | 0.95 | 0.71 |
| Butyrate (mmol/L) | 5.24<sup>B</sup> | 7.00<sup>A</sup> | 6.92<sup>A</sup> | 6.28<sup>A</sup> | 0.315 | 0.39 | 0.45 | 0.04 |
| Total VFA (mmol/L) | 88.72 | 80.92 | 75.98 | 76.55 | 3.154 | 0.63 | 0.94 | 0.39 |
| A:P | 1.80<sup>B</sup> | 1.53<sup>B</sup> | 3.74<sup>A</sup> | 3.59<sup>A</sup> | 0.174 | <0.01 | 0.51 | 0.70 |
| Ammonia (mg dL<sup>-1</sup>) | 4.06<sup>B</sup> | 8.04<sup>A</sup> | 5.78<sup>B</sup> | 8.91<sup>A</sup> | 0.543 | 0.51 | 0.02 | 0.65 |

Means with different superscripts in a row differ at $P \leq 0.05$ .

### Changes in CP concentrations exert minimal influence on rumen microbial composition in lambs

To investigate the effects of CP concentrations on the rumen microbiome, we comparatively analyzed the microbial abundance profiles under varying CP and forage concentrations using species-resolved metagenomics. Despite anticipated differences in VFA and ammonia profiles across different dietary treatments, CP concentrations (10% vs. 13%) had a minimal impact on overall rumen microbial composition, with only one species, *Butyrivibrio sp9001041551*, exhibiting an approximately 2-fold change in relative abundance between the two CP concentrations (Fig. 3a). In contrast, forage concentrations (30% vs. 70%) led to marked shifts in microbial abundance, as exemplified by substantial abundance changes (>1,000-fold) in multiple species, such as *CAG-791 sp900320025* and *Tractidigestivibacter sp900119625* (Fig. 3b and Table S5). These results suggest that the tested dietary forage levels exert a stronger influence on microbial abundance than the tested CP concentrations.

**Fig. 3:**
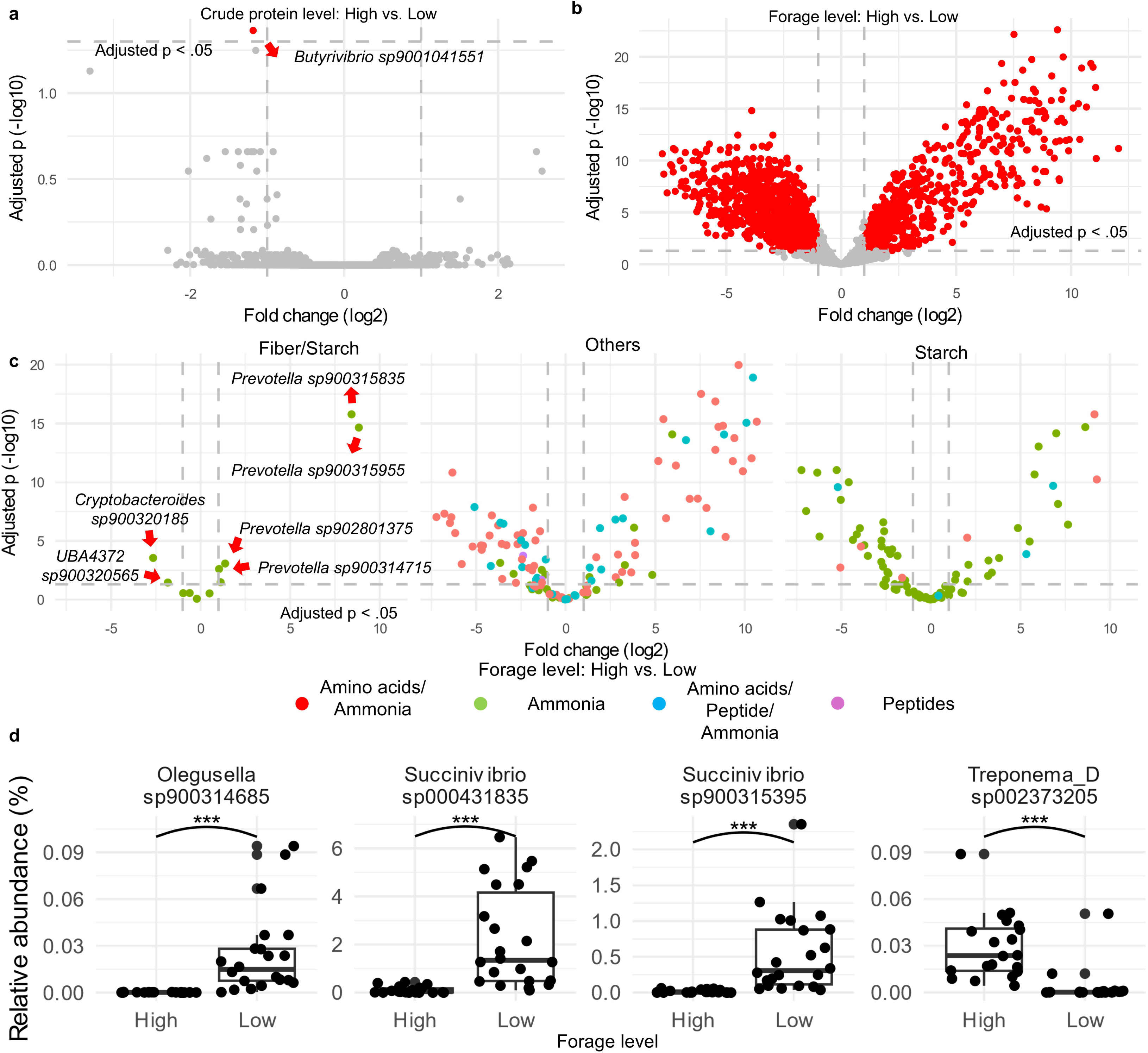
Rumen bacterial species differentially abundant in lambs fed diets with varying levels of forage and protein. **a**, log_2_-fold changes of rumen bacterial species in lambs fed low vs. high crude protein diets; **b**, Log_2_-fold changes of rumen bacterial species in lambs fed low vs. high forage diets; **c.** log_2_-fold change of rumen bacterial species with varying polysaccharide and nitrogen utilization capacities in lambs fed low vs. high forage diets; **d**, relative abundance of rumen ureolytic bacteria differentially abundant in lambs fed diets with different forage levels; **e**, relative abundance of rumen bacterial species exhibiting a significant interaction effect between forage and crude protein levels. Diets: LFLP, low forage with low crude protein; LFHP, low forage with high crude protein; HFLP, high forage with low crude protein; and HFHP, high forage with high crude protein.

Building on the substrate utilization capacities inferred from our comparative genomic analysis (Table S4), we further analyzed microbial relative abundance based on their substrate utilization capacities (Fig. 3c). Ammonia served as the primary N source among the examined rumen bacteria capable of hydrolyzing both plant cell wall and starch (as evidenced by genes encoding both cellulase and amylase). While most bacteria displayed only marginal change in relative abundance, *Prevotella* sp900315835 and *Prevotella* sp900315955 exhibited substantial increases in relative abundance (approximately 100-fold) in the high forage groups (Fig. 3c). In contrast, bacteria lacking the ability to hydrolyze both starch and fiber demonstrated versatile N utilization strategies, with species differing in N utilization capacities showing either decreased or increased relative abundance. Similarly, bacteria capable of hydrolyzing starch but not fiber predominantly utilize ammonia as their sole N source, and they displayed variable shifts in relative abundance across dietary treatments. Collectively, forage rather than CP concentrate plays a more critical role in shaping the abundance of key microbial groups with specific N utilization capacities.

Of the ureolytic bacteria identified previously, four species were detected (Fig. 3d). *Treponema_D sp002373205* was significantly more predominant in the high-forage group, while the other three species were significantly more predominant in the low-forage group. Overall, differential abundance analyses revealed that the tested CP concentrations (10 vs. 13%) had minimal effect on the relative abundance of most rumen bacteria, regardless of their N utilization strategies. Notably, only three bacterial species exhibited CP-forage concentrations interactions (Fig. S4), a pattern also observed at the genus level (Fig. S5). These findings align with previous work reporting similar rumen microbial composition between calves fed high vs. low CP (12.5% vs. 17.3) diets [66], where improved NUE in calves fed a high-CP diet was attributed to adequate net energy supplies. However, an *in vitro* study demonstrated that soluble dietary protein availability—not total CP—shaped the rumen microbial composition [67], underscoring the importance of soluble dietary protein availability in driving shifts within the rumen. Collectively, the CP treatments in this study did not appear to significantly affect the microbial composition, whereas forage levels and soluble protein availability are more significant drivers of rumen microbiome structure and NUE.

### At the community level, gene expression profiles vary depending on fermentation profiles

To understand the relationship between dietary treatments and N assimilation, we examined the metatranscriptomic profiles across the four dietary treatments (Fig. 4). Of the N assimilation-related genes, only *nad*X (encoding aspartate dehydrogenase) showed differential expression across the dietary treatments, Specifically, *nad*X expression was significantly elevated in the low forage groups relative to the other dietary treatments (Fig. 4a). The lack of variation further corroborates the findings that many rumen bacteria possess limited regulatory responses to fluctuations in rumen ammonia concentration. Interestingly, *gdh*A (encoding glutamate dehydrogenase) consistently showed the highest expression across dietary treatments, followed by *gln*A (encoding glutamine synthetase) and *ans*B (encoding asparagine synthetase). Notably, the *nad*X expression was comparable to that of *gdh*A in the low-forage groups, while in the high-forage groups, the *nad*X expression was similar to that of *gln*A and *ans*B. These findings align with an earlier enzymatic study that revealed high glutamate dehydrogenase and aspartate dehydrogenase activities in sheep fed different diets (whole barley, whole barley supplemented with 3% urea, or dried grass combined with concentrate) [46]. However, with a more pronounced difference in dietary concentrate concentrations, our study revealed a significant shift in aspartate dehydrogenase expression that does not align with the previous enzymatic assay. Moreover, the negligible activity of alanine dehydrogenase detected enzymatically [46] aligns with the low expression levels of *ald* (alanine dehydrogenase) and *asp*A (aspartate ammonia-lyase/aspartase). However, the minimal glutamine synthetase activity reported in the enzymatic assays [46] contrasts with the high gene expression level of this enzyme observed in this study. Because many rumen bacteria, including the dominant *Prevotella* species, encode a complete GS-GOGAT pathway (Fig. 1), the limited glutamine synthetase activity detected in the enzymatic assays might be attributed to experimental limitations, including isoenzyme variability and interference from endogenous components (such as the presence of other ATP-utilizing enzymes). Additionally, variability at the translational and post-translational levels might also cause discrapencies between enzymatic activity and gene expression. These discrapencies underline the challenges in directly linking gene expression to enzymatic activity and emphasize the need for multi-omic analyses, including metaproteomic anslysis. Collectively, our findings suggest that diet-induced variability in N assimilation can be mediated by transcriptional and post-transcriptional mechanisms, particularly for key enzymes such as aspartate and glutamate dehydrogenases.

**Fig. 4:**
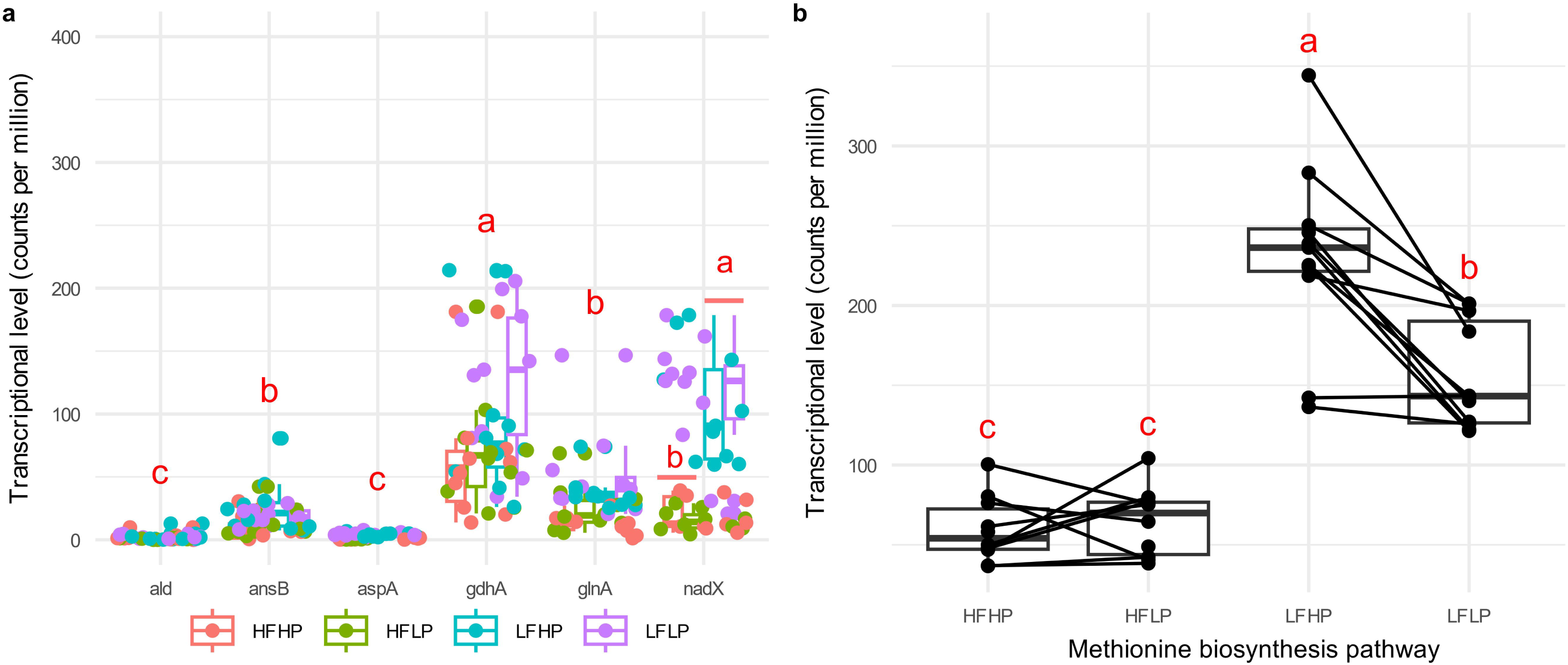
Transcriptional levels of nitrogen assimilation-related genes (a) and pathways (b) significantly differing across dietary treatments. Treatments with different letters (a, b, and c) differ significantly (*P* < 0.05). Transcriptional levels were determined at the reads level using HUMAnN3. Diets: LFLP, low forage with low crude protein; LFHP, low forage with high crude protein; HFLP, high forage with low crude protein; and HFHP, high forage with high crude protein.

Optimization of dietary amino acid profiles—particularly through targeted supplementation with rumen-protected essential amino acids like methionine, lysine, and histidine—improves NUE in ruminants. However, these effects were non-significant when diets contained adequate metabolizable protein [68–70]. To mechanistically investigate this further, we examined gene expression at the level of metabolic pathways, focusing on individual amino acid biosynthesis pathways under different dietary treatments. Using the HUMAnN3 pipeline, we found no significant differential expressions of most pathways except for the methionine biosynthesis pathway (EC.2.3.1.31, EC.2.5.1.49, and EC.2.1.1.13/EC.2.1.1.14). This pathway exhibited the highest expression in the low-forage-high-protein group, followed by the low-forage-low-protein group, while lambs fed the high-forage diets showed the lowest expression irrespective of CP concentrations (Fig. 4b). Supplementation with rumen-protected methionine rhas been demonstrated to improve NUE in dairy cows fed diets deficient in metabolizable protein, including elevated N conversion (milk N/N intake) and lowered N excretion [68, 70]. These results suggest that methionine biosynthesis is tightly regulated by dietary protein and forage levels, with important implications for optimizing amino acid supplementation strategies to enhance NUE in ruminants.

### Rumen bacteria exhibit diverse gene expression responses accompanied by shifts in fermentation profiles

To explore gene expression responses of rumen bacteria to varying dietary conditions, we further examined the genome-resolved gene expression profiles, focusing on the 37 prevalent MAGs detected in more than 80% of the samples at either forage level (Fig. 5a). Among these species, four species—*Intestinibaculum porci*, *Succinivibrio sp000431835*, *RUG023 sp900315435*, *UBA2810 sp900317945*—showed both high relative abundance and transcriptional activity, with all being more abundant in the low-forage groups. In addition, *I. porci* was also more abundant in the low-forage-high-protein group compared to the low-forage-low-protein group (Fig. 5b). Among N assimilation-related genes, *nad*X was found exclusively in *RUG023 sp900315435* (Fig. S6). Further genomic analyses revealed that *I. porci* and *Succinivibrio sp000431835* were ureolytic, and their urease expression levels did not differ between the low- and high-protein groups irrespective of the forage level (Fig. 5c). Genes encoding key methionine biosynthesis enzymes, including O-acetylhomoserine (thiol)-lyase and O-acetylhomoserine/O-acetylserine sulfhydrylase were expressed at higher levels in *UBA2810 sp900317945* in the high-protein-low-forage group than in the low-protein-low-forage group (Fig. 5d and 5e). In contrast, the other two genes of the methionine biosynthesis pathway (i.e., methionine synthase and 5-methyltetrahydrofolate-homocysteine methyltransferase) showed no differences in expression (Fig. S7). Since *UBA2810 sp900317945* exhibited the highest overall expression, the significant increase in methionine biosynthesis pathway expression at the community level was probably driven predominantly by this species. Notably, methionine is also essential for polyamine biosynthesis, and bacteria likely recycle methionine continuously to maintain a steady supply for polyamine synthesis and other methylation processes, thereby downregulating its *de novo* synthesis. This premise is supported by the observation that a bolus dosing of methionine reduced its degradation while simultaneously increasing extracellular methionine accumulation [71]. Thus, the increased expression of methionine biosynthesis genes in *UBA2810 sp900317945* likely represents a strategy to balance salvage with biosynthetic flexibility, ensuring metabolic efficiency in the high-protein, low-forage rumen environment. Additionally, it may reflect a diet-dependent lag in D-methionine racemization at the community level, although further investigation is needed to verify this hypothesis.

**Fig. 5:**
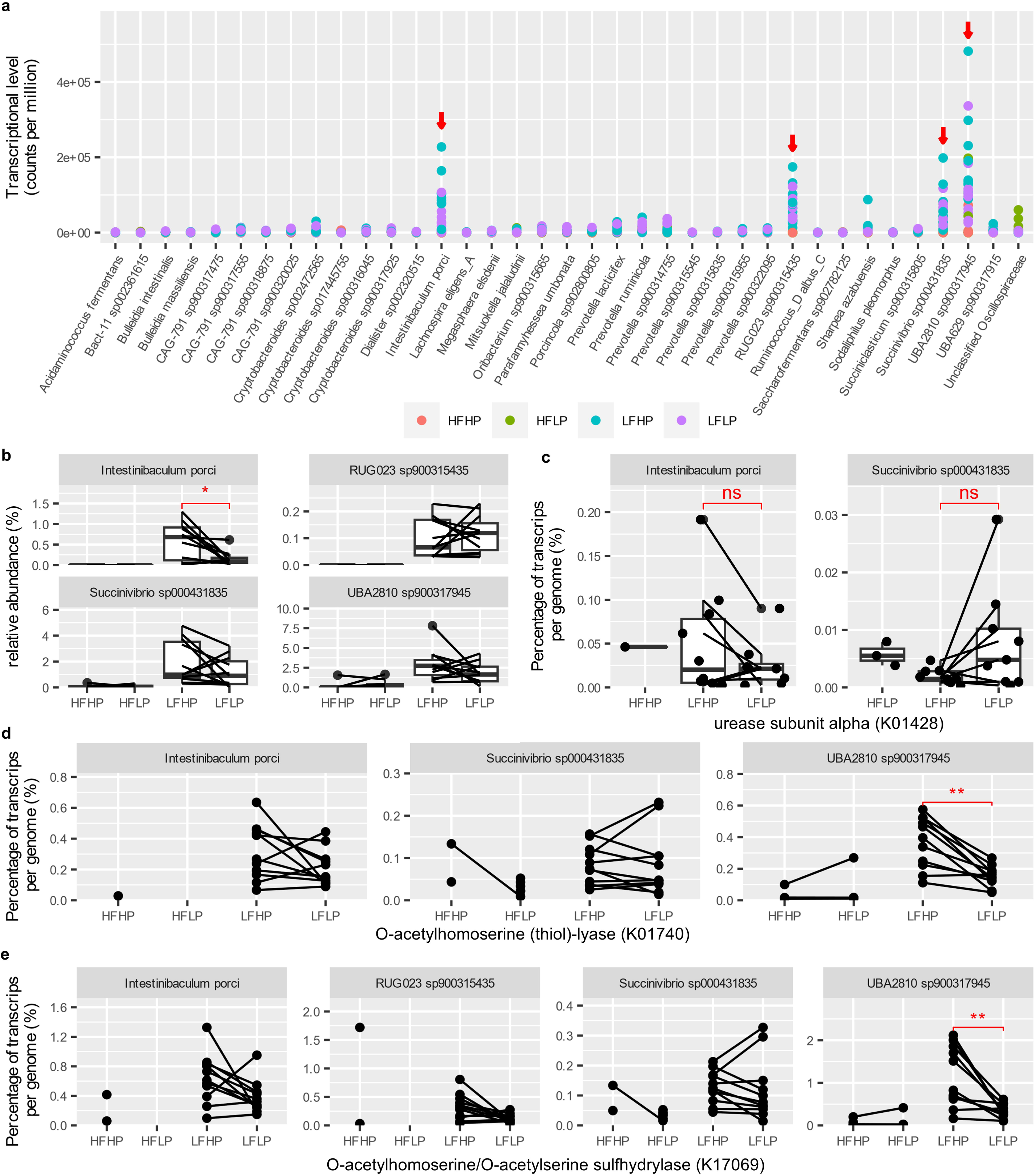
Gene expression profiles of prevalent bacterial species in lambs fed diets with varying levels of forage and protein. **a**, Overall transcriptional levels of rumen bacteria that exhibited high transcriptional activities. The red arrows indicate four species that exhibited exceptional transcriptional activities; **b**, relative abundance of transcripts of the four species; **c**, transcriptional level of urease alpha subunit in two species with high transcriptional activities; **d**, transcriptional level of O-acetylhomoserine (thiol)-lyase genes involved in the methionine biosynthesis pathway in three species exhibiting high transcriptional activities; **e**, transcriptional level of O-acetylhomoserine/O-acetylserine sulfhydrylase in highly transcriptional active species. Diets: LFLP, low forage with low crude protein; LFHP, low forage with high crude protein; HFLP, high forage with low crude protein; and HFHP, high forage with high crude protein.

The lack of differential expression in N assimilation-related genes, coupled with elevated expression of the methionine biosynthesis pathway, suggests that rumen bacteria adapt to varying ammonia availability by modulating *de novo* amino acid synthesis rather than altering N assimilation, This premise is consistent with a previous study on *E. coli*, which showed similar growth rates when utilizing aspartate or ammonia as the sole N source [72]. Furthermore, carbohydrate fermentability likely influences amino acid biosynthesis by affecting the availability of ATP and the pool of metabolic intermediates (e.g., pyruvate, oxaloacetate, 3-phosphoglycerate, and alpha-ketoglutarate), as indicated by the elevated expression of methionine biosynthesis pathways in *UBA2810 sp900317945* in the low-forage groups. Optimizing carbohydrate fermentability in diets could enhance microbial amino acid synthesis indirectly, presenting a practical approach to improving dietary NUE.

Peptidases are integral to N metabolism in the rumen ecosystem. To investigate their roles, we analyzed the expression profiles of total peptidases (Fig. 6a) and specific peptidase families (Fig. 6b) encoded by species exhibiting high gene expression. Despite variations in dietary composition, no differences were observed in the overall peptidase expression levels. However, each species expressed distinct peptidase families tailored to specialized strategies for polysaccharide degradation and N utilization (Table S4 and Fig. S6). This observation underscores the functional diversity among rumen bacteria and their capability to coordinate dietary protein degradation with carbohydrate utilization. These findings suggest that microbial specialization likely enhances overall nutrient utilization in the rumen, which is critical for optimizing dietary NUE. Additionally, rumen protozoa harbor a wide array of peptidases [73] and profoundly influence intra-ruminal N recycling and NUE [10], yet the specific functions of their peptidases and their relationship to NUE remain underexplored. Systematic characterization of protozoal peptidases could reveal novel pathways in ruminal N metabolism, with direct implications for enhancing feed efficiency and productivity in ruminants.

**Fig. 6:**
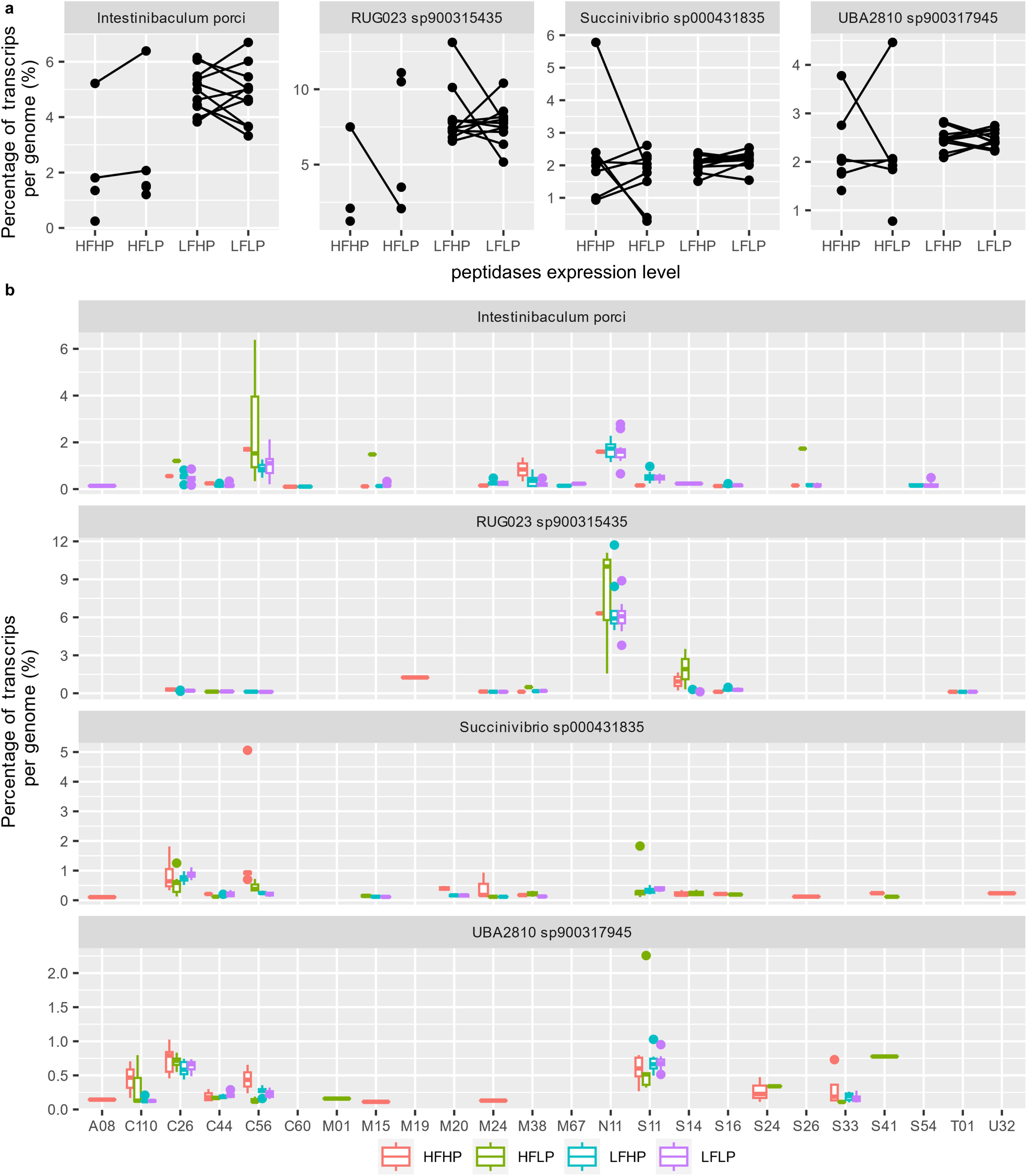
Peptidase gene expression profiles of species with high transcriptional activities in lambs fed diets with varying levels of forage and protein. **a**, transcriptional levels of total peptidases; **b**, the transcriptional level of each peptidase family encoded by each transcriptionally highly active species. Diets: LFLP, low forage with low crude protein; LFHP, low forage with high crude protein; HFLP, high forage with low crude protein; and HFHP, high forage with high crude protein.

To achieve improved NUE, ruminant nutritionists have investigated various strategies, including synchronizing N and energy availability and optimizing the amino acid profiles of rumen-degradable protein [6, 69, 74, 75]. However, the outcomes have been inconsistent or of limited practical significance, likely due to, at least partially, the complexity of rumen fermentation and its regulatory mechanisms. In this study, using comparative genomics and genome-resolved multi-omics analysis, we provide mechanistic insights into the N utilization strategies of diverse rumen bacteria, which are both compositionally and transcriptionally predominant under varying dietary conditions. Overall, our findings corroborate previous animal trials and enzymatic assays at both community and genome levels, but the species-specific N utilization strategies identified in this study underscore the complexity of N assimilation and the necessity of genome-resolved investigations to better understand and optimize N utilization for improved ruminant productivity. Furthermore, this study advances our understanding of rumen microbial ecology and highlights opportunities to develop nutritional strategies targeting species-specific metabolic pathways.

## DATA AVAILABILITY

All sequencing data and the MAGs assembled in this study have been deposited in the NCBI database with the accession ID PRJNAXXXXX.

## Acknowledgments

This work is supported in part by the USDA National Institute of Food and Agriculture (Award number: 2021-67015-33393). We thank Sripoorna Somadundaram and Ming Weng of Dr. Yu’s lab for their assistance in rumen sample collection. We also thank the Ohio Supercomputer Center for providing the computing resources.

## AUTHOR CONTRIBUTIONS

MY, ZY, JF, and AR conceived the project. MY and AR collected the samples and carried out the experiments. MY performed bioinformatic analyses. JG manually curated the phage genome annotations. MY and ZY wrote the manuscript. All authors revised the manuscript. All authors have read and approved the final manuscript.

---

**Fig. S1: Schematic representation of the animal trial.**

**Fig. S2: Overview of nitrogen assimilation regulatory genes carried by 192 rumen bacterial species.**

Outer rings (in red, from innermost to outermost) represent the presence of genes encoding leucine-responsive regulatory protein, glutamine synthetase adenylyltransferase, PII uridylyl-transferase, glutamine synthetase repressor, two-component system NtrB/NtrC (also NRII/NRI), PII protein, and nitrogen assimilation regulatory protein. Each leaf represents one species, and its color represents the phylum to which the species belongs. The branch within the same color belongs to the same genus (white or grey). Each species is represented by at least 2 genomes. A gene (or gene cluster) is considered present in a species only when at least 80% of the genomes within that species carry this gene (or gene cluster). A gene (or gene cluster) is considered absent in a species only when the gene is absent in all of the genomes within the species (or gene cluster).

**Fig. S3: Overview of nitrogen utilization genes carried by rumen archaea.** Each species is represented by at least 2 genomes. A gene (or gene cluster) is considered present in a species only when at least 80% of the genomes within that species carry this gene (or gene cluster). A gene (or gene cluster) is considered absent in a species only when the gene is absent in all of the genomes within the species (or gene cluster).

**Fig. S4: Differential abundance of rumen bacterial species across various dietary treatments that showed a significant interaction between CP and forage levels.**

Diets: LFLP, low forage with low crude protein; LFHP, low forage with high crude protein; HFLP, high forage with low crude protein; and HFHP, high forage with high crude protein.

**Fig. S5: Rumen bacterial genera differentially abundant across the dietary treatments.**

**a**, Log_2_-fold change of rumen bacterial genera in low vs. high forage diets; **b**, log_2_-fold change of rumen bacterial genera in low vs. high crude protein diets; **c**, relative abundance of rumen microbial genera exhibiting a significant interaction between forage and crude protein levels. Diets: LFLP, low forage with low crude protein; LFHP, low forage with high crude protein; HFLP, high forage with low crude protein; and HFHP, high forage with high crude protein.

**Fig. S6: Expression levels of nitrogen assimilation genes in prevalent species in lambs fed different diets.**

Diets: LFLP, low forage with low crude protein; LFHP, low forage with high crude protein; HFLP, high forage with low crude protein; and HFHP, high forage with high crude protein.

**Fig. S7: Transcriptional levels of genes encoding methionine synthase (a) and 5-methyltetrahydrofolate--homocysteine methyltransferase (b).**

Diets: LFLP, low forage with low crude protein; LFHP, low forage with high crude protein; HFLP, high forage with low crude protein; and HFHP, high forage with high crude protein.

