## Supplementary material for "New Insights into Microbial Nitrogen Utilization in the Rumen Enabled by Genome-Resolved Multi-Omics": Table S1.docx

|  | HFLP | HFHP | LFLP | LFHP |
| --- | --- | --- | --- | --- |
| Grass hay (%) | 70.00 | 70.00 | 30.00 | 30.00 |
| Soybean hulls (%) | 6.43 | 0.00 | 20.43 | 0.00 |
| Ground corn (%) | 20.01 | 11.44 | 45.01 | 48.44 |
| DDGS (%) | 0.00 | 15.00 | 1.00 | 18.00 |
| Limestone (%) | 1.00 | 1.00 | 1.00 | 1.00 |
| Trace mineral salt (%) | 0.50 | 0.50 | 0.50 | 0.50 |
| Vitamin A (%) | 0.01 | 0.01 | 0.01 | 0.01 |
| Vitamin D (%) | 0.01 | 0.01 | 0.01 | 0.01 |
| Vitamin E (%) | 0.05 | 0.05 | 0.05 | 0.05 |
| Vitavet Selenium (%) | 0.09 | 0.09 | 0.09 | 0.09 |
| AV fat blend (%) | 1.50 | 1.50 | 1.50 | 1.50 |
| Ammonium Chloride (%) | 0.40 | 0.40 | 0.40 | 0.40 |
| CP (%) | 10.27 | 13.18 | 10.31 | 13.12 |

**Table S1:** Composition of diets containing high or low forage (HF or LF, respectively) and high or low protein (HP or LP, respectively)
