## Supplementary material for "New Insights into Microbial Nitrogen Utilization in the Rumen Enabled by Genome-Resolved Multi-Omics": Table S3.docx

| Carbohydrate source | Nitrogen source | Number of species (number of genera) |
| --- | --- | --- |
| Fiber (cellulose + hemicellulose) | Amino acids + ammonia | 2 (2) |
|  | Peptides + amino acids + ammonia | 1 (1) |
| Starch | Amino acids | 18 (15) |
|  | Ammonia | 157 (36) |
|  | Peptides + amino acids + ammonia | 5 (4) |
| Fiber (cellulose + hemicellulose) + starch | Amino acids | 1 (1) |
|  | Ammonia | 21 (10) |
| Others | Amino acids | 143 (87) |
|  | Ammonia | 48 (28) |
|  | Peptides + amino acids + ammonia | 53 (40) |

**Table S3: The polysaccharide hydrolysis and nitrogen assimilation capacities of rumen microbes illustrated in Fig. 1.**
