## Supplementary material for "New Insights into Microbial Nitrogen Utilization in the Rumen Enabled by Genome-Resolved Multi-Omics": Table S5.docx

Table S5. Detailed results of the bacterial species shown in Fig. 3c.

| log2-Fold  Change (High vs. Low forage) | Species | Carbon source | Nitrogen source |
| --- | --- | --- | --- |
| -1.43 | *Prevotella sp902776665* | starch | ammonia |
| 4.86 | *Prevotella sp900315545* | starch | ammonia |
| 3.81 | *Prevotella sp900315525* | starch | ammonia |
| 1.41 | *Prevotella sp002354095* | starch | ammonia |
| 1.36 | *Prevotella sp900315035* | starch | ammonia |
| 3.40 | *Prevotella sp002394385* | starch | ammonia |
| -2.37 | *Prevotella sp902769705* | starch | ammonia |
| 2.10 | *Prevotella sp002353225* | starch | ammonia |
| 5.51 | *Prevotella sp900314755* | starch | ammonia |
| 1.37 | *Prevotella sp902801375* | misc | ammonia |
| 2.05 | *Prevotella sp900315095* | starch | ammonia |
| -2.67 | *Prevotella sp900314935* | starch | ammonia |
| 1.12 | *Prevotella sp902779455* | misc | ammonia |
| 1.03 | *Prevotella sp900314715* | misc | ammonia |
| 6.02 | *Prevotella sp902792635* | starch | ammonia |
| 8.43 | *Prevotella sp900315835* | misc | ammonia |
| 7.66 | *Prevotella sp900316565* | starch | ammonia |
| 8.84 | *Prevotella sp900315955* | misc | ammonia |
| 5.79 | *Prevotella sp002480935* | starch | ammonia |
| 7.00 | *Prevotella sp900316015* | starch | ammonia |
| -1.83 | *UBA4372 sp900320565* | misc | ammonia |
| -4.35 | *UBA1179 sp900318995* | starch | ammonia |
| 3.22 | *Alloprevotella sp900316575* | starch | ammonia |
| -6.22 | *Cryptobacteroides sp900316675* | starch | ammonia |
| -7.23 | *Cryptobacteroides sp017445755* | starch | ammonia |
| -5.26 | *Cryptobacteroides sp900319685* | starch | ammonia |
| -6.97 | *Cryptobacteroides sp902768665* | starch | ammonia |
| -2.27 | *Cryptobacteroides sp902801885* | starch | ammonia |
| -5.03 | *Cryptobacteroides sp902761655* | starch | ammonia |
| -4.13 | *Cryptobacteroides sp900321655* | starch | ammonia |
| -2.61 | *Cryptobacteroides sp902780935* | starch | ammonia |
| -2.74 | *Cryptobacteroides sp900320965* | starch | ammonia |
| -3.71 | *Cryptobacteroides sp002393895* | starch | ammonia |
| -2.65 | *Cryptobacteroides sp900320185* | misc | ammonia |
| -2.76 | *Cryptobacteroides sp902792815* | starch | ammonia |
| -2.40 | *Cryptobacteroides sp900318565* | starch | ammonia |
| -3.96 | *Cryptobacteroides sp017427385* | starch | ammonia |
| -2.65 | *Cryptobacteroides sp902779785* | starch | ammonia |
| 7.10 | *Cryptobacteroides sp002472565* | starch | ammonia |
| -4.58 | *Cryptobacteroides sp900321245* | starch | ammonia |
| -1.86 | *Cryptobacteroides sp902785825* | starch | ammonia |
| -2.15 | *Cryptobacteroides sp900318955* | starch | ammonia |
| -6.34 | *Bact-11 sp002361615* | starch | ammonia |
| -3.52 | *Bact-11 sp902778855* | starch | ammonia |
| 1.33 | *Egerieousia sp016285455* | other | ammonia |
| -2.91 | *UBA1711 sp900317125* | other | ammonia |
| -1.61 | *UBA1711 sp902790215* | other | ammonia |
| 4.83 | *Phil12 sp900314725* | other | ammonia |
| -1.67 | *Sodaliphilus sp902762385* | starch | ammonia |
| 8.63 | *Sodaliphilus pleomorphus* | starch | ammonia |
| -2.42 | *F23-D06 sp017527945* | starch | ammonia |
| -2.51 | *RUG11690 sp902771655* | starch | ammonia |
| -2.27 | *Limivicinus sp902791215* | other | misc_n |
| 5.45 | *RUG678 sp900321335* | other | amino acids |
| -2.02 | *Ruminococcus flavefaciens_Q* | starch | ammonia |
| 2.92 | *Ruminococcus flavefaciens_R* | other | amino acids |
| 3.88 | *Ruminococcus sp002394695* | other | amino acids |
| 3.30 | *Ruminococcus flavefaciens_V* | other | ammonia |
| -4.12 | *Ruminiclostridium_E sp902780695* | other | amino acids |
| -6.39 | *Ruminococcus_E sp900315085* | other | amino acids |
| 2.79 | *Ruminococcus_E sp002350765* | other | amino acids |
| -4.25 | *Ruminococcus_E sp902776375* | other | amino acids |
| -2.69 | *Ruminococcus_E sp900315605* | other | amino acids |
| -1.87 | *Ruminococcus_E sp902797225* | other | amino acids |
| -3.31 | *Ruminococcus_E sp900317315* | other | amino acids |
| -3.77 | *UBA1213 sp900317795* | other | amino acids |
| -1.34 | *UBA1213 sp902785555* | other | ammonia |
| -5.10 | *Acetatifactor sp017527205* | other | misc_n |
| -2.78 | *Eubacterium_Q sp900321215* | other | amino acids |
| -4.18 | *Butyrivibrio fibrisolvens_C* | other | misc_n |
| -1.43 | *Butyrivibrio sp000423945* | other | amino acids |
| -2.69 | *Butyrivibrio sp000424285* | other | amino acids |
| -1.83 | *Butyrivibrio sp900104155* | other | amino acids |
| -5.20 | *Pseudobutyrivibrio ruminis* | starch | misc_n |
| -2.77 | *CAG-411 sp902795895* | starch | ammonia |
| 3.19 | *CAG-791 sp900319245* | other | misc_n |
| 8.08 | *CAG-791 sp900315055* | other | misc_n |
| 10.46 | *CAG-791 sp900320025* | other | misc_n |
| 10.09 | *CAG-791 sp900317475* | other | misc_n |
| -3.91 | *UBA1258 sp900100895* | starch | amino acids |
| -1.60 | *RUG11200 sp902766825* | starch | amino acids |
| -1.69 | *NK4A136 sp900102065* | other | amino acids |
| -1.50 | *RUG191 sp015057165* | other | amino acids |
| -1.76 | *RUG191 sp002373675* | other | amino acids |
| -2.39 | *NK4A144 sp000621405* | other | amino acids |
| -4.73 | *NK4A144 sp902779445* | other | amino acids |
| 8.84 | *Oribacterium sp900315665* | other | misc_n |
| -6.17 | *CAG-632 sp902783585* | other | amino acids |
| 6.83 | *Agathobacter faecis* | starch | misc_n |
| -1.14 | *Eubacterium_F xylanophilum* | other | ammonia |
| -1.12 | *RGIG4930 sp017470735* | other | misc_n |
| 2.03 | *UBA2821 sp002351535* | starch | amino acids |
| 1.91 | *Weimeria sp900100095* | other | misc_n |
| 3.29 | *Weimeria sp900315505* | other | amino acids |
| 1.98 | *RUG306 sp900316075* | other | misc_n |
| 1.34 | *Lachnobacterium bovis* | other | misc_n |
| 2.78 | *Lachnobacterium sp900113385* | other | misc_n |
| 5.17 | *UBA629 sp900316665* | other | amino acids |
| 6.71 | *RUG760 sp900315265* | other | misc_n |
| -3.48 | *UBA2862 sp017512905* | other | misc_n |
| -2.49 | *UBA2862 sp902765305* | other | misc_n |
| -4.69 | *RUG563 sp905236495* | other | amino acids |
| -3.65 | *UBA2943 sp002350845* | other | misc_n |
| -1.59 | *Ventricola sp002395225* | other | misc_n |
| -1.37 | *RUG472 sp902798825* | other | misc_n |
| -2.38 | *UBA3857 sp017509785* | other | peptide |
| -7.22 | *Saccharofermentans sp902782125* | other | amino acids |
| -5.81 | *Saccharofermentans sp002450095* | other | amino acids |
| -5.20 | *Saccharofermentans sp900314745* | other | amino acids |
| -6.46 | *Saccharofermentans sp902764825* | other | amino acids |
| -4.62 | *Saccharofermentans sp900176545* | other | amino acids |
| -1.35 | *Firm-16 sp900317035* | other | peptide |
| -2.44 | *Firm-16 sp017545245* | other | amino acids |
| -2.43 | *RUG11894 sp902773665* | other | misc_n |
| -1.66 | *RUG11894 sp017626845* | other | misc_n |
| 5.95 | *Hornefia butyriciproducens* | other | ammonia |
| 6.15 | *Eubacterium_T pyruvativorans* | other | amino acids |
| 5.60 | *Ornithomonoglobus sp902796445* | other | amino acids |
| 8.78 | *Pseudoramibacter fermentans* | other | amino acids |
| 3.66 | *Succinivibrio dextrinosolvens_A* | other | amino acids |
| 7.37 | *Succinivibrio sp900315395* | other | amino acids |
| 8.93 | *UBA2810 sp900317945* | other | amino acids |
| 3.82 | *UBA2804 sp900319705* | other | ammonia |
| 1.70 | *RGIG3394 sp017446355* | other | amino acids |
| -6.77 | *Anaerovibrio lipolyticus* | other | amino acids |
| -3.57 | *Selenomonas_B ruminantium_A* | other | amino acids |
| -1.33 | *Selenomonas_B ruminantium_B* | other | amino acids |
| 7.86 | *Selenomonas_C bovis* | other | amino acids |
| 9.42 | *UBA2913 sp002349925* | other | amino acids |
| -2.06 | *Succiniclasticum sp002342505* | other | amino acids |
| 9.65 | *Acidaminococcus fermentans* | other | amino acids |
| 7.56 | *Acidaminococcus sp900315205* | other | amino acids |
| 9.89 | *Megasphaera elsdenii* | other | amino acids |
| 8.36 | *Caecibacter massiliensis* | other | amino acids |
| 10.38 | *Dialister succinatiphilus* | other | amino acids |
| 2.98 | *Enteromonas sp900316365* | other | ammonia |
| 6.92 | *UBA4951 sp900320745* | other | amino acids |
| 3.87 | *Enterosoma sp900313465* | other | amino acids |
| 8.36 | *Absicoccus porci* | other | amino acids |
| 9.27 | *Intestinibaculum porci* | starch | amino acids |
| 5.33 | *Intestinibaculum sp900317015* | starch | misc_n |
| -3.66 | *Anaeroplasma sp017524905* | other | ammonia |
| 9.33 | *RUG023 sp900315435* | other | amino acids |
| -6.33 | *CADAEX01 sp902787125* | other | amino acids |
| -5.05 | *UBA3636 sp902793555* | starch | amino acids |
| 1.44 | *UBA1367 sp902779675* | other | misc_n |
| 10.67 | *Tractidigestivibacter sp900119625* | other | amino acids |
| 8.53 | *UBA7748 sp900314535* | other | amino acids |
| 3.20 | *UBA6984 sp900321735* | starch | ammonia |
| 3.15 | *Fibrobacter sp001603905* | other | amino acids |
| -3.54 | *Fibrobacter elongatus* | other | amino acids |
| 9.14 | *Desulfovibrio sp900319575* | starch | amino acids |

misc, both fiber and starch; misc_n, ammonia and amino acids/peptides.
