## Supplementary figures and images for "New Insights into Microbial Nitrogen Utilization in the Rumen Enabled by Genome-Resolved Multi-Omics"

### Fig. S1.jpg

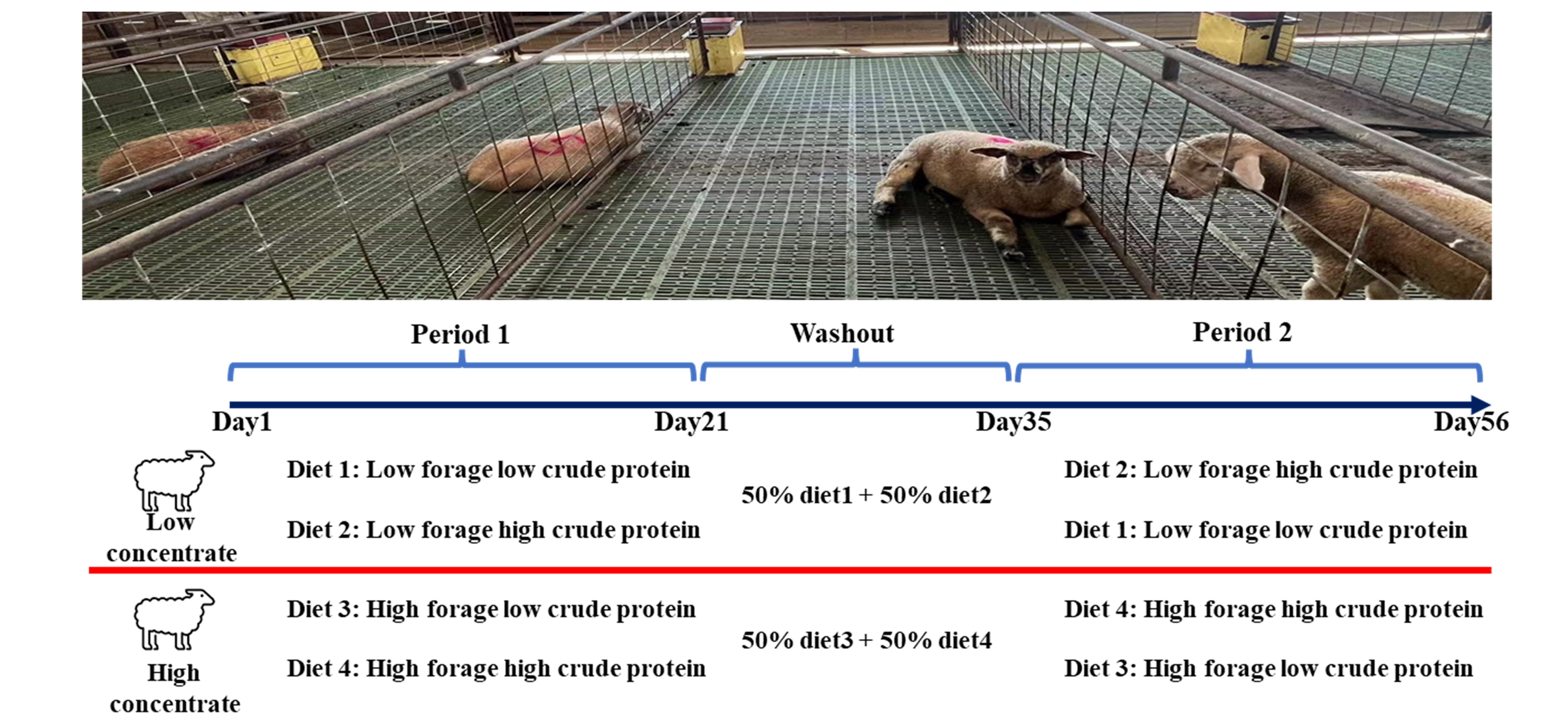

### Fig. S2.jpg

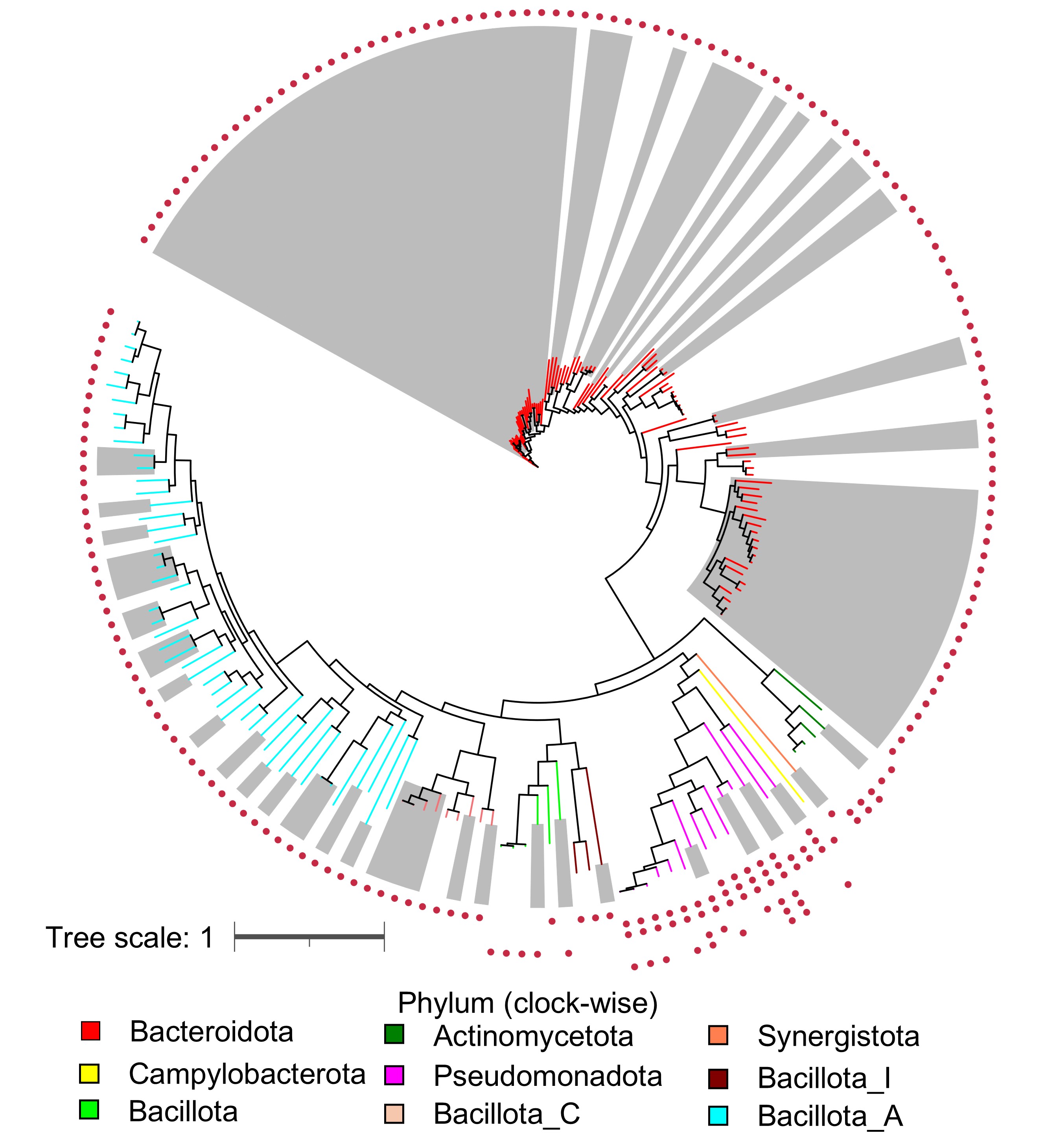

### Fig. S3.jpg

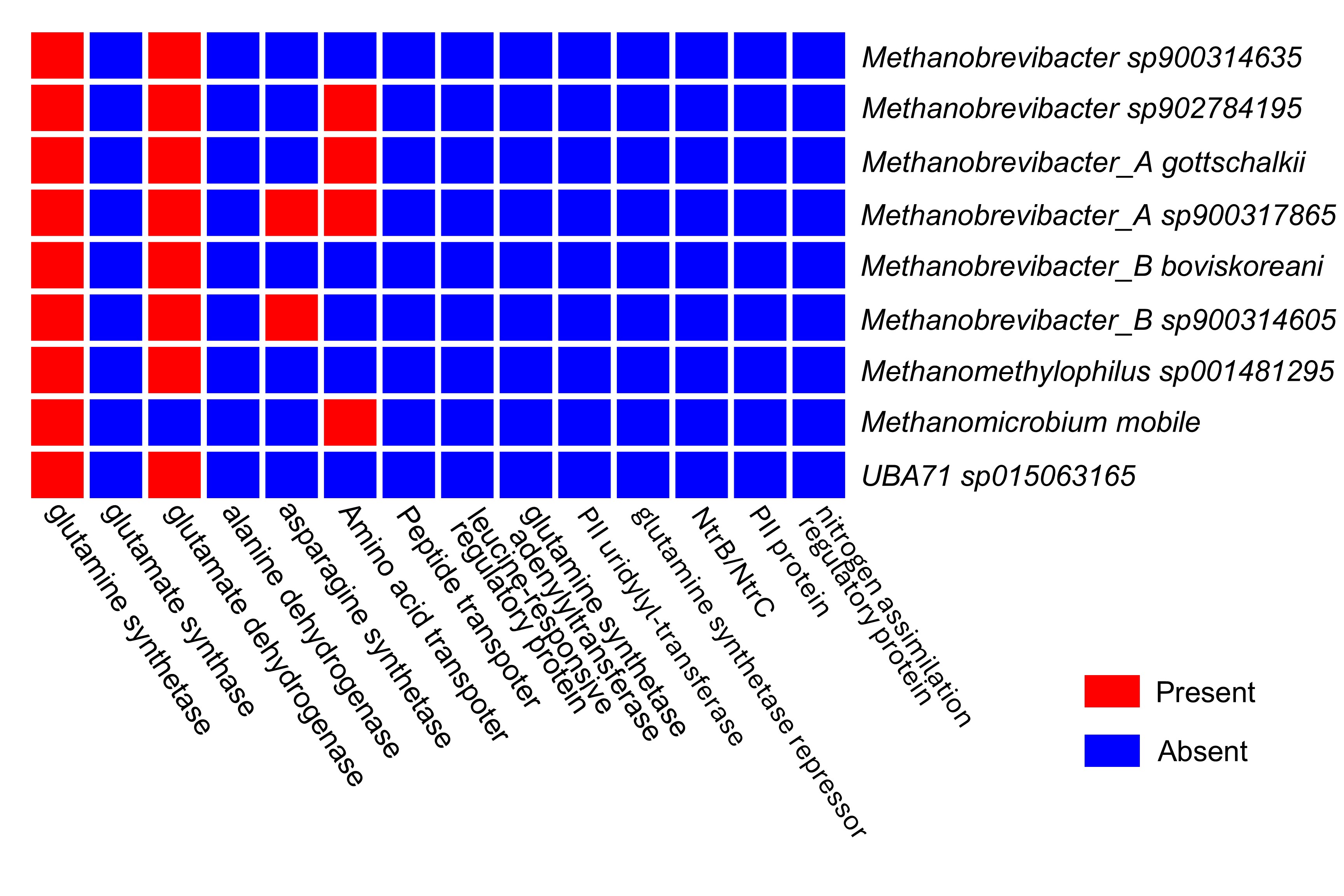

### Fig. S4.jpg

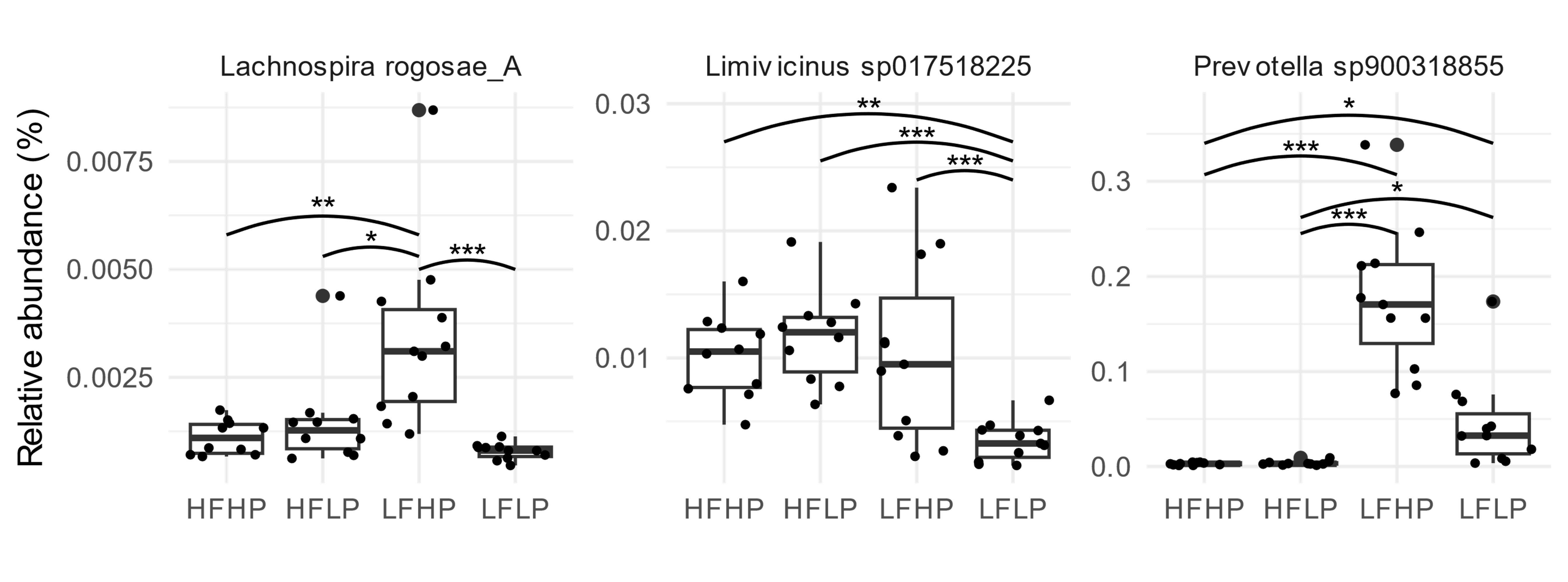

### Fig. S5.jpg

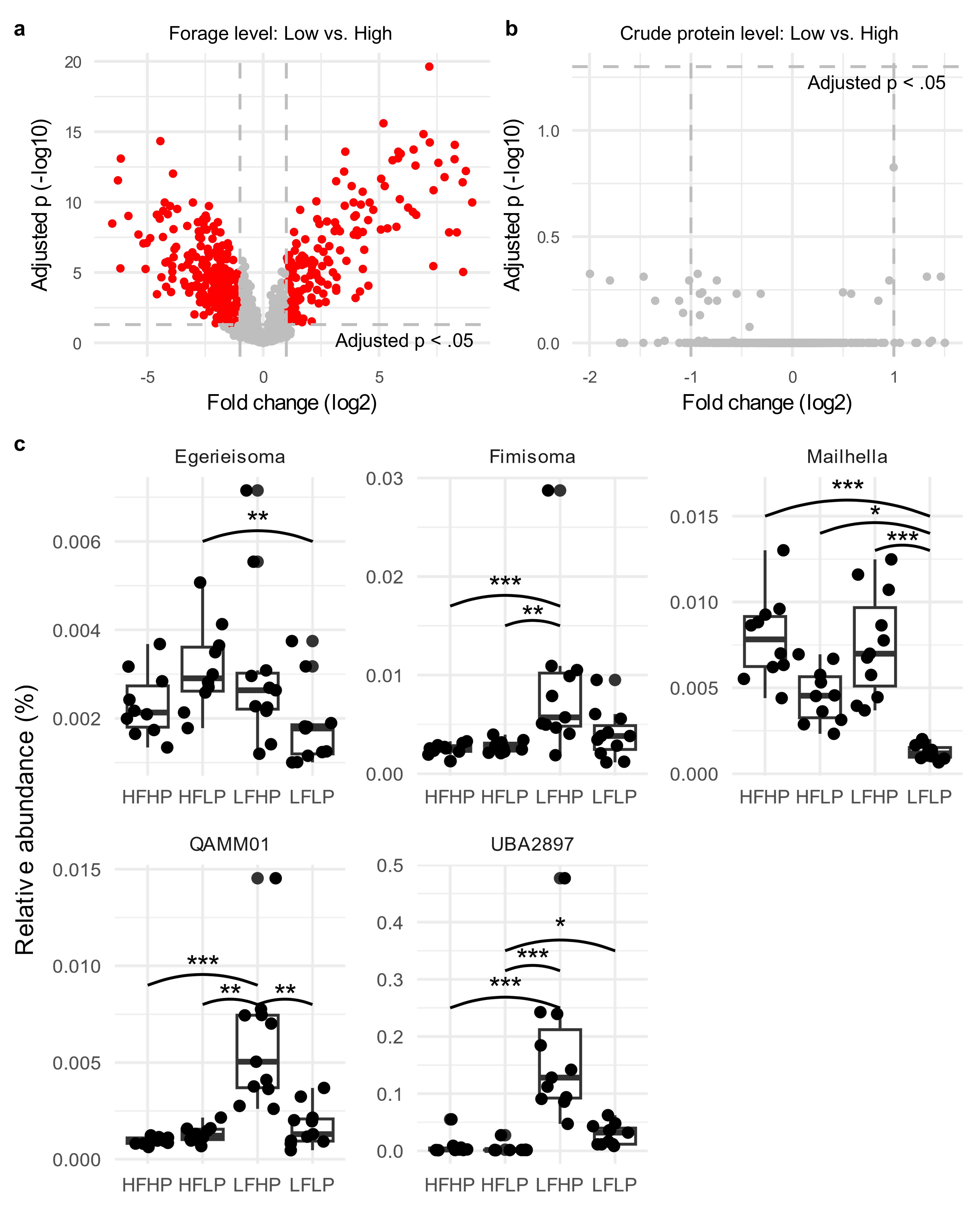

### Fig. S6.jpg

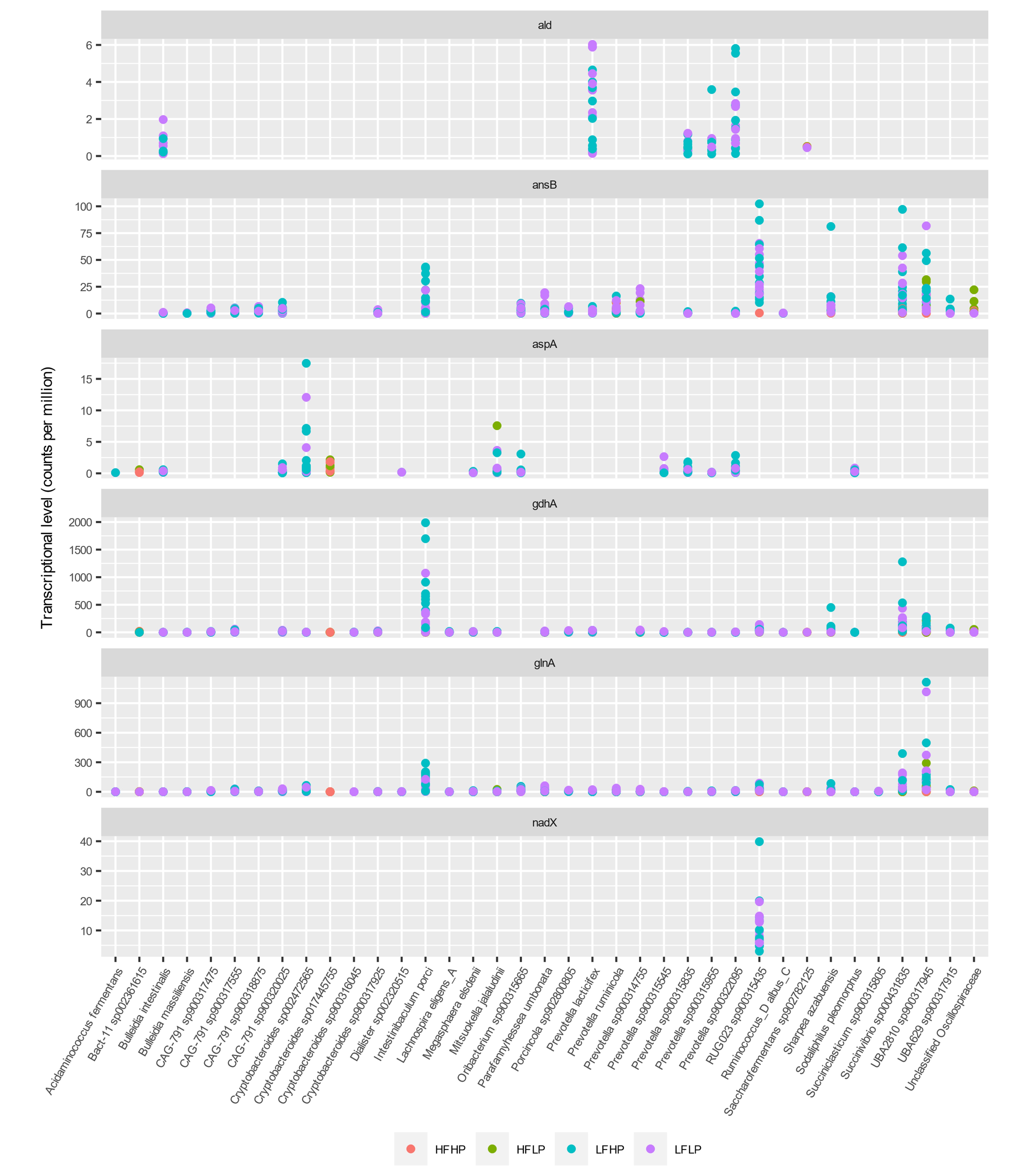

### Fig. S7.jpg

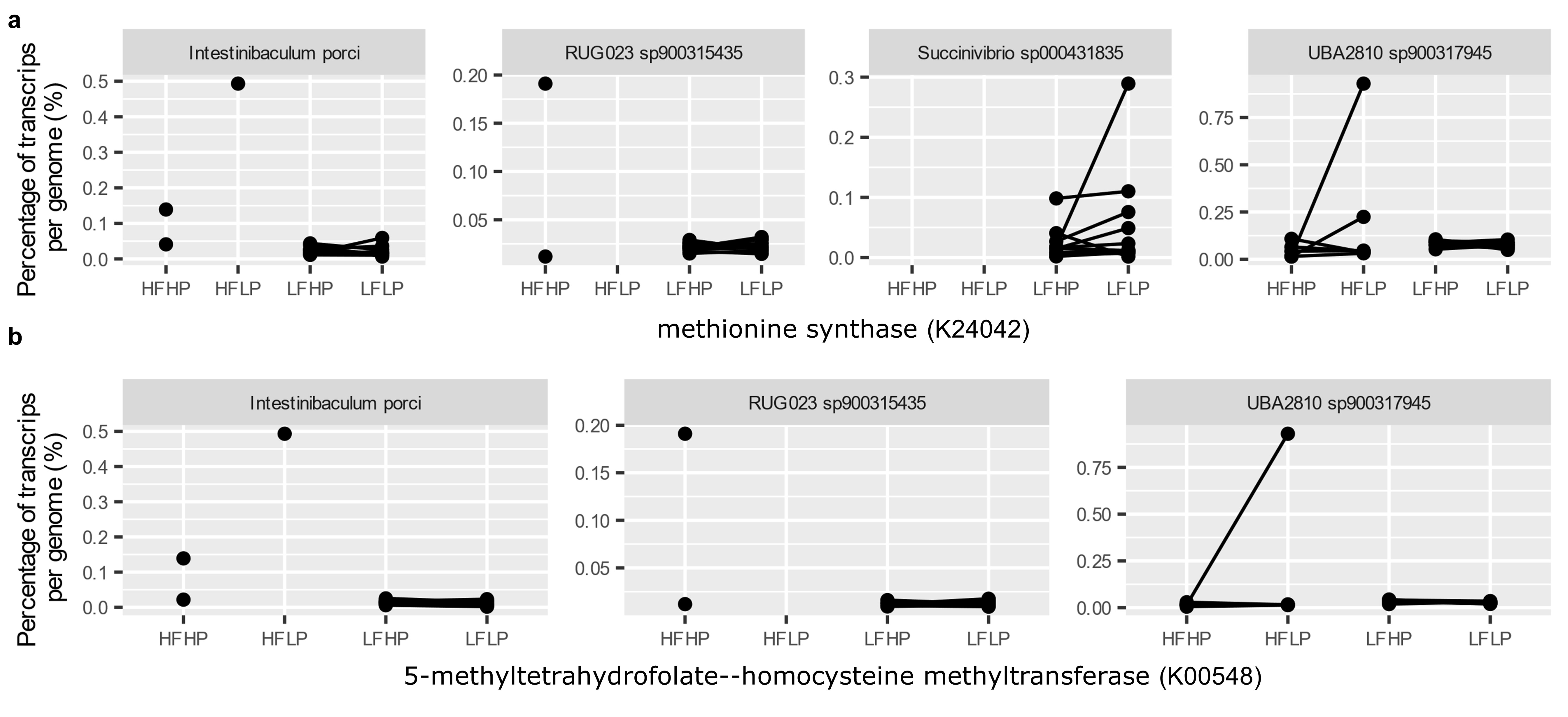
